# Microglia-Dependent and Independent Modulation of Brain Lipid Metabolism in Alzheimer’s Disease Revealed by Pharmacological and Genetic Microglial Depletion

**DOI:** 10.1101/2024.11.18.624173

**Authors:** Ziying Xu, Sepideh Kiani Shabestari, Savannah Barannikov, Kevin F. Bieniek, Mathew Blurton-Jones, Juan Pablo Palavicini, Xianlin Han

**Affiliations:** Sam and Ann Barshop Institute for Longevity and Aging Studies, University of Texas Health Science Center San Antonio, TX 78299, USA; Department of Neurobiology & Behavior, University of California - Irvine, Irvine, CA 92697, USA; Glenn Biggs Institute for Alzheimer’s and Neurodegenerative Diseases, University of Texas Health Science Center San Antonio, TX 78299, USA; Department of Pathology and Laboratory Science, University of Texas Health Science Center San Antonio, TX 78299, USA; Institute for Memory Impairments and Neurological Disorders, University of California - Irvine, Irvine, CA 92697, USA; Department of Medicine, University of Texas Health Science Center San Antonio, TX 78299, USA

## Abstract

Abnormal lipid metabolism in Alzheimer’s disease (AD) was first documented by Alois Alzheimer in his early observations of patients with a then unrecognized brain disease, which he noted was characterized by a significant presence of “adipose inclusions” or “lipoid granules”. Despite this early recognition, until recently the significance of abnormal lipid metabolism in AD has been largely overlooked by the scientific community for decades, highlighting a critical gap in our understanding of this complex disease. In the past decade, numerous loci and genes with genome-wide significant evidence of affecting AD risk have been reported. Notably, a significant portion of these AD risk genes are either preferentially or exclusively expressed by microglia in the brain and/or code for enzymes that directly or indirectly regulate lipid metabolism. This suggests a major, yet uncharacterized, role of microglia in modulating brain lipid metabolism under AD pathological conditions. In our study, we dissected microglia-dependent and independent regulation of lipid metabolism in an AD-like mouse model of amyloidosis, 5xFAD, taking advantage of pharmacological and genetic interventions to eliminate microglia. Using multidimensional mass spectrometry-based shotgun lipidomics (MDMS-SL), we identified overt changes in a number of AD-associated lipids (ADALs) in postmortem patient brains and mouse models of amyloidosis. This included bis(monoacylglycerol)phosphate (BMP), a lipid class enriched in endosomal/lysosomal compartments, and the two most abundant classes of lysophospholipids: lysophosphatidylcholine (LPC) and lysophosphatidylethanolamine (LPE), which are commonly associated with inflammation. Our findings revealed that microglial depletion prevented the accumulation of arachidonic acid-containing BMP species, which are associated with lysosomal activation induced by amyloidosis via a mechanism that involves progranulin, coded by AD risk gene *GRN*, as shown by targeted transcriptomics, immunoblotting, and immunofluorescence. Surprisingly, AD-associated LPC and LPE accumulation was not driven by microglia. Instead, LPC accumulation correlated with astrocytic activation, while LPE accumulation seems to be associated with oxidative stress. In summary, we uncovered novel microglia-dependent and independent mechanisms that drive lipid dysregulation in AD. These findings may be mechanistically linked with the early glial lipoid deposits described by Dr. Alzheimer.

Graphical Abstract.

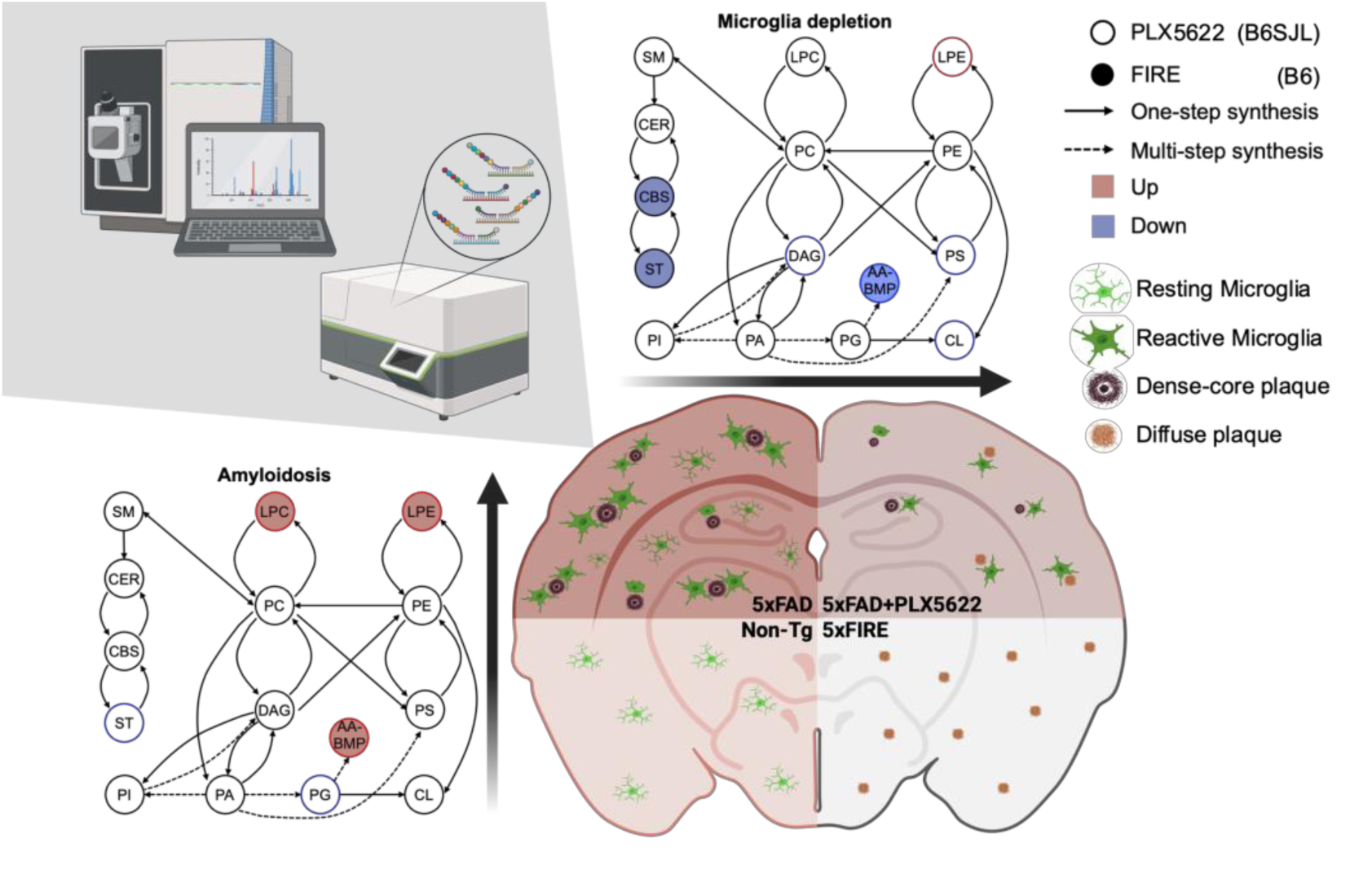

## Background/Introduction

Alzheimer’s disease (AD), the most common neurodegenerative disease and the leading cause of dementia^1^, has been characterized by impaired homeostasis in the brain and is accompanied by dramatic changes to the cellular characterization and brain microenvironment^2,3^. After being sidelined for years, the importance of the brain’s lipid metabolism disruptions is gaining increased attention by the AD research community. Amyloid-beta (Aβ) accumulation is one of the two major pathological hallmarks of AD. The accumulation of Aβ in the brain occurs early in the development of AD^4^. Recent studies suggest a significant interaction between amyloidosis and lipid metabolism. The dynamic lipid composition regulates the trafficking and activity of key membrane-bound proteins that control Aβ levels and facilitate its aggregation^5–7^. Alterations in lipid homeostasis can affect membrane fluidity, leading to a disruption in cellular signaling and increased vulnerability to amyloid plaque formation^8^. With the help of modern imaging and analysis methods, researchers underscored shifts in the metabolism of phospholipids^9^, sphingolipids^10^, cholesterol^11^, and other myelin lipids^12^ in AD development and modulating Aβ production and clearance^13^. Dysregulation of these lipids may contribute to the pathogenesis of AD, making lipid metabolism a potential target for therapeutic intervention^8^. The relationship between amyloidosis and lipid metabolism in AD underscores the complexity of the disease and suggests that a multifaceted approach, which includes the management of lipid levels, could be beneficial in slowing the progression of Alzheimer’s disease. In this study, we used two 5xFAD transgenic mouse lines (B6SJL and B6 genetic backgrounds), which recapitulate major features of Alzheimer’s disease amyloid pathology, to study the impact of amyloidosis on brain lipid metabolism.

A number of the 76 currently validated loci with genome-wide significant evidence of affecting AD risk^14^ are preferentially expressed by microglia in the brain and code for enzymes that directly or indirectly regulate lipid metabolism^15^, including *TREM2*^16,17^, *PLCG2*^18^, *INPP5D*, *GRN*^19^, among others. Microglia are known to be activated by Aβ and participate in Aβ clearance^20^. Yet long-term microglial activation can lead to reduced phagocytotic activity and increased chronic neuroinflammation^21^, which may further exacerbate AD pathologies. Manipulating colony stimulating factor 1 receptor (CSF1R) signaling, the essential signaling pathway for microglial differentiation, proliferation, and survival^22–24^, to regulate microglial population has been a hot research focus recently. The role of microglia in AD pathologies has been studied extensively in various animal models by either pharmacological elimination or deletion of microglial genes^22–24^. A series of studies have been focused on eliminating microglia by targeting the CSF1R signaling^22–24^, which is known to regulate microglial survival. Yet the effects of microglial elimination on AD pathologies are quite complex as they depend on the timing and extent of microglia depletion^25^. Multiple studies have consistently shown the CSF1R inhibitors, PLX3397 and PLX5622, by partially depleting microglia, can reduce dense core amyloid plaque load^26,27^, increase diffusible amyloid, reduce tau pathologies^28^, reduce synaptic and neuronal loss, and improve cognition^29^. However, there are also a few studies that have reported no changes in AD pathologies following partial microglial elimination^25^. On the other hand, a recent study^30^ reported that fully eliminating microglia via genetic deletion of microglia-associated *fms* intronic regulatory element (FIRE) enhancer of *Csf1r* locus in a 5xFAD background, leads to brain calcification, cerebral hemorrhages, and even premature lethality. Triggering receptor expressed on myeloid cells 2 (*TREM2*) deficiency ameliorates amyloid pathology at an early stage but exacerbates the disease progression at a late stage in the APPPS1–21 mouse model^31^. Yet, despite all the research emphasis, the effects of microglial elimination on brain lipid metabolism have been largely overlooked, except for one study specifically focused on leukotrienes, which demonstrated microglia are essential to leukotriene synthesis^32^.

Traditionally, lysophospholipids are considered key pro-inflammatory lipid markers that are found to be accumulated in neurodegeneration diseases, including AD^8^. The production of lysophospholipids is through the cleavage of membrane phospholipids, including the two most abundant ones: phosphatidylcholine (PC) and phosphatidylethanolamine (PE). Both PC and PE contain a hydrophilic head group and two hydrophobic acyl chains at sn-1 and sn-2 positions, which can be cleaved upon phospholipase activation or reactive oxygen species (ROS) attack^33,34^. The cleavage products, lysophosphatidylcholine (LPC) and lysophosphatidylethanolamine (LPE), along with their cleavage enzyme cytosolic phospholipase A_2_ (cPLA2) have been found to accumulate in AD subjects and mouse models^8,35^.

Previous studies have also shown there is an abnormal accumulation of Bis(monoacylglycero)phosphate (BMP) in neurodegenerative diseases, including Lewy body dementia^36^ and AD^6,37^. BMP, also known as lysobisphosphatidic acid (LBPA), is a unique class of phospholipids playing a vital role in cellular lipid management. It functions primarily in the late endosomes/lysosomes, where it promotes lipid organization by activating lipid hydrolases and lipid transfer proteins^38–42^. While the presence of BMP is crucial for certain cellular functions, its excessive accumulation can be a sign of underlying metabolic stresses and could contribute to disease pathogenesis. Elevated levels of BMP have been observed in lysosomal storage disorders and under energy imbalance conditions^43,44^. Additionally, an increase in BMP levels has been linked to disorders like Niemann-Pick type C disease, where it is implicated in cholesterol trafficking defects^45^.

In this study, we aimed to decipher microglia-dependent and independent mechanism(s) underlying abnormal lipid metabolism in 5xFAD by depleting microglia following both pharmacological and genetic interventions. With highly accurate multi-dimensional mass spectrometry-based shotgun lipidomics (MDMS-SL)^46^, we are able to detect several bioactive lipid classes/species abnormally altered under amyloidosis conditions. LPC, LPE, and BMP accumulate in the brains of two separate cohorts of 5xFAD mice in different genetic backgrounds. After depleting microglia, amyloidosis-induced accumulation of arachidonic acid-containing BMP (AA-BMP) is completely prevented through a mechanism that involves progranulin, while, surprisingly, LPC and LPE accumulation is unaltered or further exacerbated. To the best of our knowledge, this study is the first to describe microglia-dependent and independent regulation of global brain lipid metabolism under AD-like amyloidosis conditions, shedding novel mechanistic insights into which cell types drive specific lipid abnormalities in the AD brain.

## Results

### Abnormal lipid metabolisms in amyloidosis and microglial depletion mouse brains

Previous studies have shown the complex relationship between Aβ and lipid metabolism^5–13^. Here we used 5xFAD mice as a model of amyloidosis to study the impact of human Aβ accumulation and aggregation on the brain lipidome. Taking advantage of recent advances on microglial survival studies based on modulating CSF1R signaling, we used both a pharmacological intervention by supplementing PLX5622 orally in the diet that leads to partial microglial depletion in 5xFAD mice and a genetic intervention using the *Csf1r* enhancer deletion (FIRE) mouse model where microglia are completely eliminated to test if partial or full microglia depletion rescues the disruption of brain lipid metabolism induced by amyloidosis.

A previous study reported that PLX5622 dietary supplements (1,200 ppm) for 10 weeks were able to reduce dense-core plaques within cortical regions in 5xFAD mice^27^. PLX5622 treatment with 1,200 ppm for 24 weeks was able to reach 97–100% reduction of microglia without causing any obvious negative effects in mouse behaviors or motor function. Since this pharmacological intervention cannot fully deplete microglia, as an alternative to investigate the long-term consequences of microglial full absence, a more recent study^30^ utilized a novel mouse model that harbors a deletion of the FIRE enhancer within the *Csf1r* locus. These FIRE mice lack microglia, but have no impact on peripheral macrophages and retain normal brain anatomy and health^24^. Crossing FIRE with 5xFAD mice also reduced dense-core plaques; however, it also induced severe cerebral amyloid angiopathy, brain calcification, cerebral hemorrhages, and leading to premature mortality^24^. These conflicting results urge the need to evaluate the impact of microglia on whole-brain homeostasis in more detail. The fact that multiple microglia-specific/enriched AD risk genes are highly associated with lipid metabolism^15^ suggests the important role of microglia in regulating brain lipid metabolism in the context of AD.

To address this knowledge gap, we use both pharmacological and genetic approaches to better understand how microglia affect brain lipid metabolism under disease-like amyloidosis conditions taking advantage of MDMS-SL. 5xFAD (B6SJL) female mice were fed with 1,200 ppm PLX5622-containing OpenStandard diet (OSD, Research diets) or control diet (OSD) from 1.5 months to 5 months of age (**Fig. 1A**). FIRE male and female mice were crossed with 5xFAD (B6) mice to generate the microglia-free 5xFIRE mice (**Fig. 1B**).

**Figure 1.**
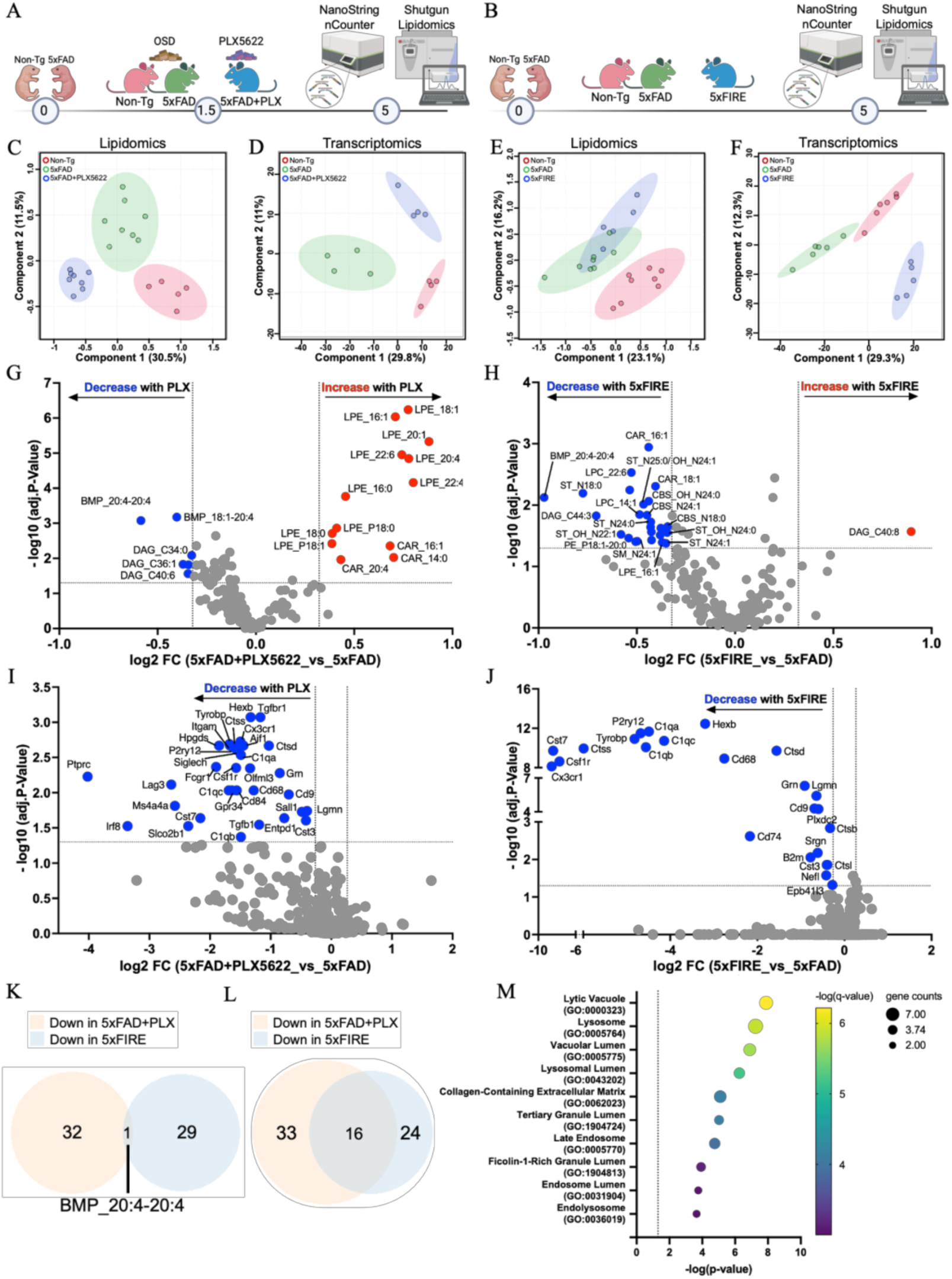
Lipidomics and targeted transcriptomics reveal abnormal metabolism in microglia-depleted AD-like mouse brains. (A) 5xFAD mice were fed with OpenStandard Diet (OSD) or OSD plus 1200 ppm PLX5622 from 1.5 to 5 months of age. (B) 5xFAD mice crossed with FIRE mice were harvested at 5 months of age. Brains collected from 5-month-old mice. Principal component analysis (PCA) of brain lipidomic profiles in pharmacological (C) and genetic (E) microglia-depleted mice, respectively. PCA of NanoString profiles in pharmacological (D) and genetic (F) microglia-depleted mice, respectively. Volcano plots showing differentially altered lipid species after depleting microglia pharmacologically (G) and genetically (H) under amyloidosis conditions. Differentially expressed genes after pharmacological (I) and genetic (J) microglial depletion under amyloidosis. Venn diagrams showing shared lipid species (K) and genes (L) that decreased after microglial depletion under amyloidosis. Data transformation: log_10_; data scaling: mean. Adjusted p ≤ 0.05 (Benjamini-Hochberg), log_2_ fold-change cutoff: 0.25. (M) Bubble plot showing top 10 Gene Ontology (GO) significant terms using the 16 DEGs from panel L.

Transformed and scaled lipidomics data from 198 lipid species normalized to total protein content were clustered in partial least squares discriminant analysis (PLS-DA) for both cohorts. The overall brain lipidome showed clear separations between Non-Tg and 5xFAD groups in both the pharmacological (**Fig. 1C**) and genetic (**Fig. 1E**) cohorts. The individual hits that contribute to the separation are displayed via volcano plots using unpaired one-factor analysis (**Fig. S1E-F**). Consistent in both cohorts, there were several LPC, LPE, and BMP species increased with amyloidosis revealed by Venn diagram analysis (**Fig. S1G**). After microglial depletion, the pharmacological intervention (**Fig. 1C**) showed a greater impact on brain lipidome than the genetic deletion cohort (**Fig. 1E**), compared to their corresponding amyloidosis groups (complete vs partial separation from 5xFAD in respective PLS-DAs). Specifically, pharmacological microglial elimination further increases several lysophospholipid species (**Fig. 1G**), which were not observed in the genetic depletion cohort (**Fig. 1H**). Yet, the genetic microglial depletion exerts an exclusive impact on brain lipidome by bringing down several myelin lipids (e.g., ST_N24:1, CBS_N24:1, and SM_N24:1) and energy lipids (**Fig. 1H**), which were not observed in pharmacological elimination cohort (**Fig. 1G**). From all the lipid species that were significantly increased in the brains of both cohorts of 5xFAD mice, partial or complete depletion of microglia was sufficient to prevent accumulation of one of them, i.e., arachidonic acid-containing (20:4) BMP (AA-BMP) (**Fig. 1K**).

Similarly, for the targeted transcriptomics, CNS NanoString analysis was able to capture distinguished gene signatures in amyloidosis and after microglial partial and complete depletion, resulting in complete separations between groups in both PLS-DAs (**Fig. 1D** and **1F**). The pathways that contribute to these separations are shown as heatmaps (**Fig. S1A-B**). The pathways that were upregulated in both 5xFAD cohorts were downregulated after microglial depletion in both cohorts, especially microglial and inflammasome-related pathways. Consistent trends were observed for individual gene hits significantly altered under amyloidosis and microglial depletion conditions for both pharmacological (**Fig. S1C** and **1I**) and genetic (**Fig. S1D** and **1J**) approaches. As a further validation of microglial depletion effects other than the reduced *Csf1r* shown in both volcano plots (**Fig. 1I-J**), reduced *Trem2* levels were confirmed in both cohorts (**Fig. S1J-K**). The stable APP and synaptic protein level were also confirmed by 6E10 and Homer1 westerns in the pharmacological cohort (**Fig. S1L** and **1M**). Venn diagram analysis revealed (or identified) 17 genes increased in both 5xFAD cohorts, which were functionally linked with lysosome-related pathways based on the Gene Ontology (GO) term analysis (**Fig. S1I**). Similar genes (**Fig. 1L**) and pathways (**Fig. 1M**) were decreased following partial or complete microglial depletion, implying that increases in lysosomal function were primarily driven by microglia. In alignment with previous lipidomics findings, the shared AA-BMP species found to be increased under amyloidosis and reduced after microglial depletion could be functionally linked to lysosomal pathway regulation.

### Microglial depletion prevents amyloidosis-induced lysosomal AA-BMP accumulation

The fact that AA-BMP is the only lipid species altered in both pharmacological (PLX5622) and genetic (FIRE) microglia-deficient cohorts, indicates that AA-BMP accumulates primarily in microglia and suggests it may be synthesized and regulated there. Importantly, we also found evidence of abnormal BMP accumulation in postmortem AD brains (**Fig. S2A**), which is consistent with the trend observed for total BMP levels in the 5xFAD mouse model (**Fig. S2B-C**). Almost all BMP species increased in human AD brains (**Fig. 2A**, **2D**, and **2G**), except DHA (22:6)-BMP which remained unaltered (**Fig. S2D**). These BMP alterations in human AD brains were well recapitulated in both 5xFAD cohorts, where AA (20:4) and oleic acid (18:1)-containing BMP species accumulated (**Fig. 2B-C**, **2E-F**, and **2H-I**) while DHA (22:6)-BMP remained unaltered (**Fig. S2E-F**). Human demographic information are presented in **Supplementary Table 1**.

**Figure 2.**
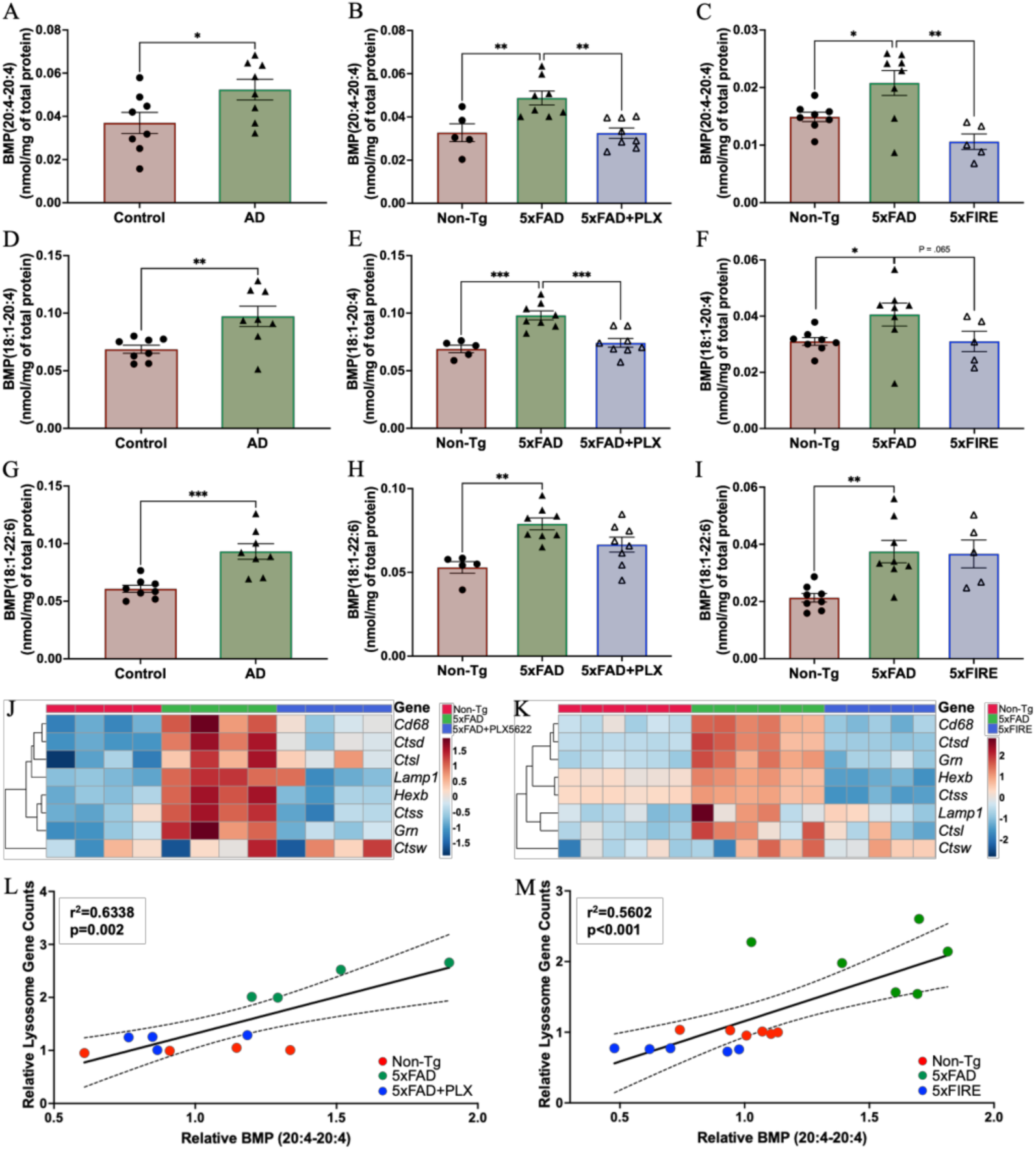
Microglial depletion rescues abnormal lysosomal BMP accumulation under amyloidosis. Specific BMP species levels in human AD brains (A, D, and G) and mouse brains where microglia were depleted pharmacologically (B, E, and H) or genetically (C, F, and I). All data presented as mean ± SEM, normalized to WT. Ordinary one-way ANOVA with Turkey correction, *p ≤ 0.05, **p ≤ 0.01, and ***p ≤ 0.001. Heatmaps of lysosomal gene changes following pharmacological (J) and genetic (K) approaches to deplete microglia. Data transformation: log10; data scaling: mean. Correlations between relative AA-BMP level and relative total lysosome gene counts in microglia-depleted mouse brains following pharmacological (L) and genetic (M) approaches.

Specific BMP species respond differently towards microglial depletion as well. AA-containing BMPs (BMP_20:4-20:4 and BMP_18:1-20:4) were decreased following pharmacological (**Fig. 2B** and **2E**) and genetic (**Fig. 2C** and **2F**) microglial depletion. Docosahexaenoic acid (DHA)-containing BMP (BMP_18:1-22:6) only shows amyloidosis-related accumulation (**Fig. 2G-I**), but not microglia-dependent reduction (**Fig. 2H-I**), while DHA-specific BMP (BMP_22:6-22:6) doesn’t seem to respond to microglial elimination since its levels are unchanged in either pharmacological (**Fig. S2E**) or genetic (**Fig. S2F**) cohorts.

Consistently, we also found abnormal upregulation of lysosome-related genes under amyloidosis of 5xFAD mice in both cohorts (**Fig. 2J-K**). Both pharmacological and genetic microglial depletion largely restored the abnormal increase of lysosomal gene expression to non-Tg levels (**Fig. 2J-K**). The microglia/macrophage-specific lysosomal marker (CD68) protein level was also found to be upregulated in 5xFAD brains (**Fig. S2G-H**). As an additional indication of lysosomal function, abnormal hyperglycosylated LAMP1 induced by amyloidosis was restored by microglial elimination (**Fig. S2G** and **S2J**), while normal LAMP1 level remained unaltered (**Fig. S2G** and **S2I**), indicating microglial elimination was able to rescue lysosomal impairments. As previously mentioned, BMPs are located in the late endosome/lysosome system, and their levels have been considered an indicator of proper lysosomal function^23^. As expected, AA-BMPs are highly correlated with NanoString lysosomal gene scores (**Fig. 2L**-**M**).

### Microglial progranulin as a putative modulator of AA-BMP

A transcriptional gene network interference study^47^ suggested that lysosomal function and organization are mediated by a secreted glycoprotein, progranulin (PGRN) which encoded by gene *granulin* (*GRN*). *GRN* mutations lead to frontotemporal dementia^48^ and amyotrophic lateral sclerosis^49^. Clinical case^19^ and GWAS^50^ studies have also linked *GRN* with AD. In line with previous studies, *Grn* expression is found to be upregulated in amyloidosis mouse models (**Fig. 3A** and **3C**), while pharmacological and genetic microglial depletion both prevented its amyloidosis-induced upregulation (**Fig. 3A** and **3C**). PGRN protein levels were also largely attenuated (**Fig. 3B)**. More recently, a study showed that progranulin knockout mice result in the loss of BMP in the brain^51^, demonstrating that progranulin regulates BMP metabolism. Consistently, progranulin content highly correlates with AA-BMP levels in both cohorts (**Fig. 3D-F**). In the brain, progranulin is highly expressed by microglia and neurons^52^. Immunofluorescence studies revealed that microglial elimination reduces microglia-specific (plaque-associated) PGRN accumulation, but in contrast, seems to promote neuronal PGRN expression (**Fig. 3G-I)**. This trend matches the observed BMP species-specific responses toward microglial elimination, with AA-containing BMPs showing microglia-dependent regulation while DHA-containing BMPs display an increasing trend (**Fig. S2F**).

**Figure 3.**
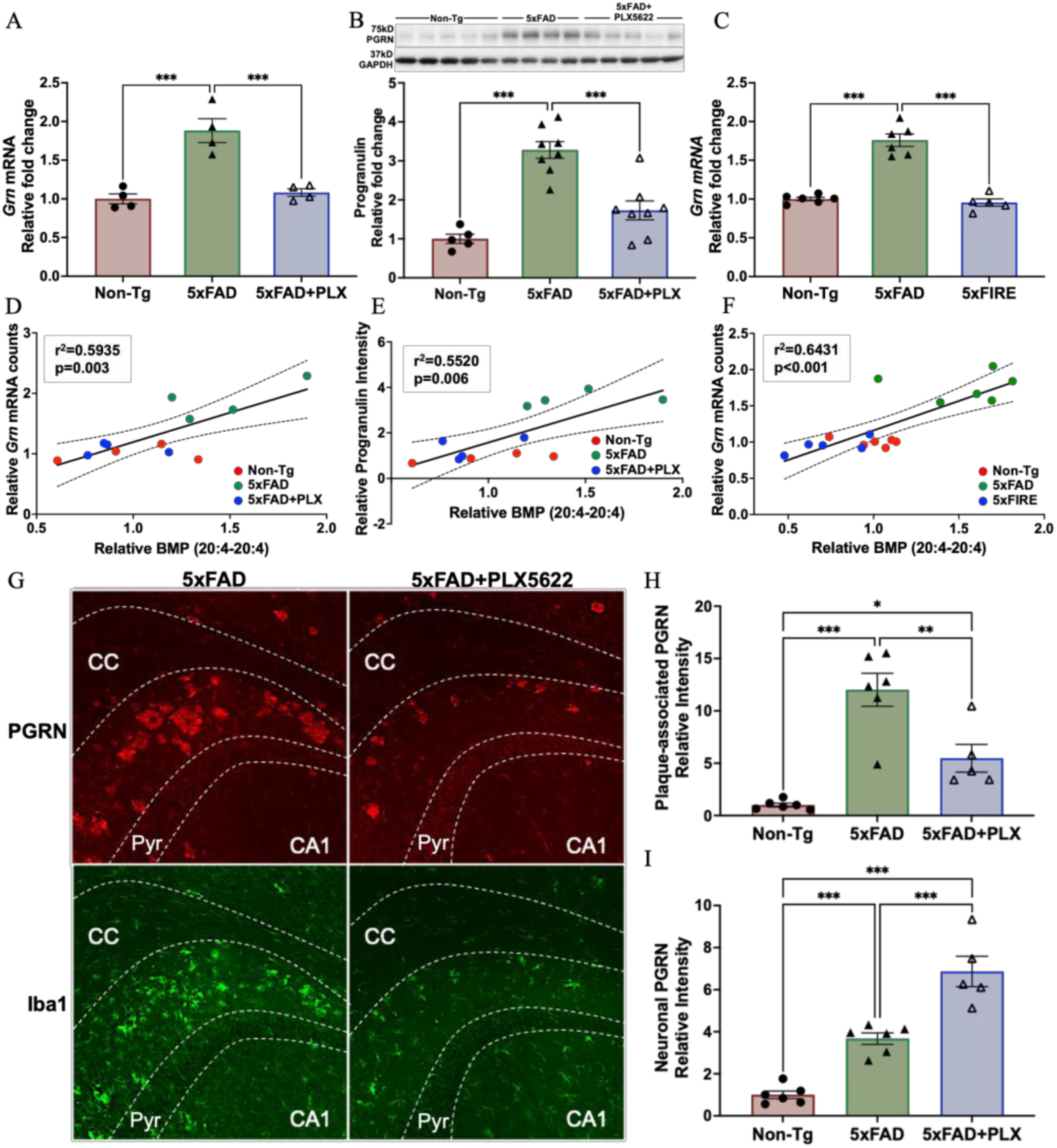
Progranulin levels increase in response to amyloid deposition in a microglia-dependent manner and correlate with arachidonic acid-containing BMP. Relative *GRN* mRNA changes in pharmacological (A) and genetic (C) microglia-depletion conditions. Correlations between relative AA-BMP levels and relative *GRN* mRNA counts in pharmacological (D) and genetic (F) microglia-deficiency conditions. Relative progranulin levels (B) and its correlation with AA-BMP content (E). Immunofluorescence (G) of microglia (Iba1^+^) and progranulin (PGRN^+^) in 5xFAD and 5xFAD+PLX5622. CC: corpus callosum; CA1: hippocampal subfields cornu ammonis; Pyr: pyramidale. Red: progranulin; green: Iba1. Quantification of plaque-associated progranulin intensity (H) and neuronal progranulin intensity (I). Quantification was done for hippocampus regions. All data presented as mean ± SEM, normalized to WT. Ordinary one-way ANOVA with Turkey correction, *p ≤ 0.05, **p ≤ 0.01, and ***p ≤ 0.001.

### Microglia do not drive abnormal amyloid-induced lysophospholipids accumulation

In addition to BMP, the two most abundant lysophospholipids, lysophosphatidylcholine (LPC) and lysophosphatidylethanolamine (LPE), are also found to be changed with amyloidosis and/or microglial depletion. Lysophospholipids have been proposed to serve as a biomarker of AD since they have been consistently shown to accumulate in human brains and almost all AD mouse models^6,34,53–56^. Accumulated lysophospholipids have been associated with neuroinflammation. However, little is known about which cell type is the major contributor to lysophospholipid production in the brain.

LPC is generated from phosphatidylcholine (a major constituent of cell membranes) upon phospholipase A_2_ cleavage. Saturated fatty acids-containing (SFA) and monounsaturated fatty acids-containing (MUFA) LPCs are believed to exert pro-inflammatory roles in promoting the secretion of chemotactic factors and the production of reactive oxygen species (ROS)^57–60^. Polyunsaturated fatty acid-containing (PUFA) LPC species, such as LPC_22:4 and LPC_22:6, are known to have anti-inflammatory effects since they help ameliorate the inflammatory impact induced by saturated LPC *in vivo*^60–62^. Consistently, in this study, total LPC levels are found to accumulate in human AD brains (**Fig. 4A**) and under amyloidosis (**Fig. 4B-C**), an effect primarily driven by the pro-inflammatory SFA and MUFA LPCs (**Fig. S4A-C**). Microglial depletion with pharmacological and genetic approaches under amyloidosis has a minor impact on the total LPC level (**Fig. 4B-C**). Interestingly, although other species remain unchanged in both cohorts after microglial depletion, there was a significant reduction of PUFA-LPC (LPC_22:6) after genetic microglial depletion (5xFIRE) (**Fig. S4C**), implying a reduction of anti-inflammatory regulation. Overall, our data demonstrate that amyloidosis induces an accumulation of pro-inflammatory LPCs in a microglia-independent manner.

**Figure 4.**
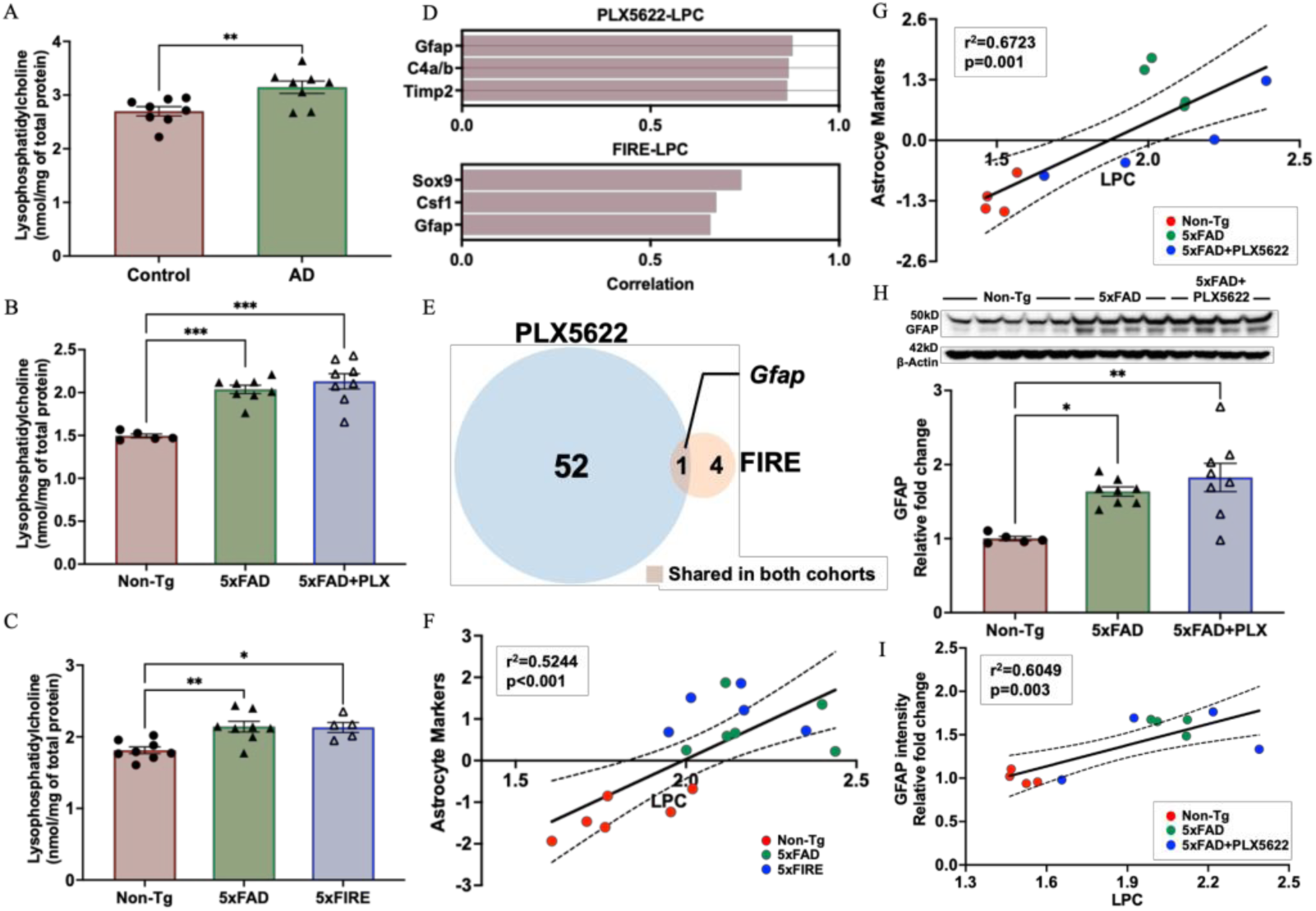
Alzheimer’s disease-associated lysophosphatidylcholine accumulation is not restored by microglial elimination, instead correlates with astrocytic markers. Total LPCs in human AD brains (A) and in pharmacological (B) and genetic (C) microglial-depleted mouse brains. Two tailed ordinary one-way ANOVA with Turkey correction, *p ≤ 0.05, **p ≤ 0.01, and ***p ≤ 0.001. (D) Top three gene hits that significantly correlate with LPCs in pharmacological and genetic cohorts. Data transformation: cube root for the pharmacological cohort, log_10_ for the genetic cohort; data scaling: mean for the pharmacological cohort, pareto for the genetic cohort. (E) Venn diagram shows the overlapping gene between two cohorts that shows correlation with LPCs. Correlation analysis between LPC and astrocyte marker pathway score in pharmacological (G) and genetic (F) cohorts. (H) Western-blot of GFAP (top) and its quantification (bottom) in the pharmacological cohort. Two tailed ordinary one-way ANOVA with Turkey correction, *p ≤ 0.05, **p ≤ 0.01, and ***p ≤ 0.001. (I) Correlation between LPC and GFAP in the pharmacological cohort. All data is presented as mean ± SEM.

To gain insights into the mechanisms underlying amyloidosis-induced LPC accumulation, we first investigated phospholipase A_2_ (PLA_2_), which cleaves PC to generate LPC. Increased PLA_2_ activity has been reported in the brains of AD patients and animal models, including a previous study from our group where we observed increased phosphorylated (activated) calcium-dependent and - independent phospholipase A_2_ activity (phospho-cPLA_2_ and iPLA_2_, respectively)^34^. Contrary to our hypothesis, neither iPLA_2_ nor phospho-cPLA_2_ was significantly altered under amyloidosis or PLX5622 treatment (**Fig. S5A-C**). We then did correlation analyses between total LPC levels and individual gene changes under different conditions. The top three positively correlated genes in both pharmacological and genetic cohorts were astrocytic (**Fig. 4D**). Based on this finding, we did further pathway correlation analyses, and found that LPC strongly correlated with astrocyte markers in both the pharmacological (**Fig. 4G**) and genetic (**Fig. 4F**) cohorts. Upon unbiased correlation screening among LPC and transcriptomic pathways from NanoString, other pathways that stood out (show significant correlations of p<0.05, R^2^>0.4) included JAK-STAT (**Fig. S3C** and **S3G**) and cytoskeletal dynamics (**Fig. S3D** and **S3H**), both related to astrocyte activation. In addition, the only gene found to be positively correlated with LPC that was shared in both cohorts is *Gfap* (**Fig. 4E**). As expected, *Gfap* (the mRNA levels shown in **Fig. S3A** and **S3E**) also correlates with LPCs in both the pharmacological (**Fig. S3B**) and genetic (**Fig. S3F**) cohorts. GFAP Western blot analysis revealed a significant increase of GFAP levels in 5xFAD that tended to be further increased in 5xFAD+PLX5622 (**Fig. 4H**). Consistent with the high correlations at the gene level, LPC is also highly associated with GFAP at the protein level (**Fig. 4I**). In summary, instead of being regulated in a microglia-dependent manner, LPC accumulation is more likely driven by astrocytes.

The other major lysophospholipid class, LPE, is produced by cleavage of phosphatidylethanolamine (a major constituent of cell inner membranes and intracellular organelle membranes). LPE has been demonstrated to regulate calcium signaling^63^, exhibit anti-apoptotic activity, and enhance neuronal differentiation and migration^64^. Importantly, LPE abnormal accumulation has also been linked with oxidative stress^65,66^. In this study, we found that total LPE levels increased consistently in human AD brains (**Fig. 5A**) and mouse models of amyloidosis (**Fig. 5B-C**), with the 18:1, 20:4, and 22:6 being the most significantly affected species (**Fig. S4D-F**).

**Figure 5.**
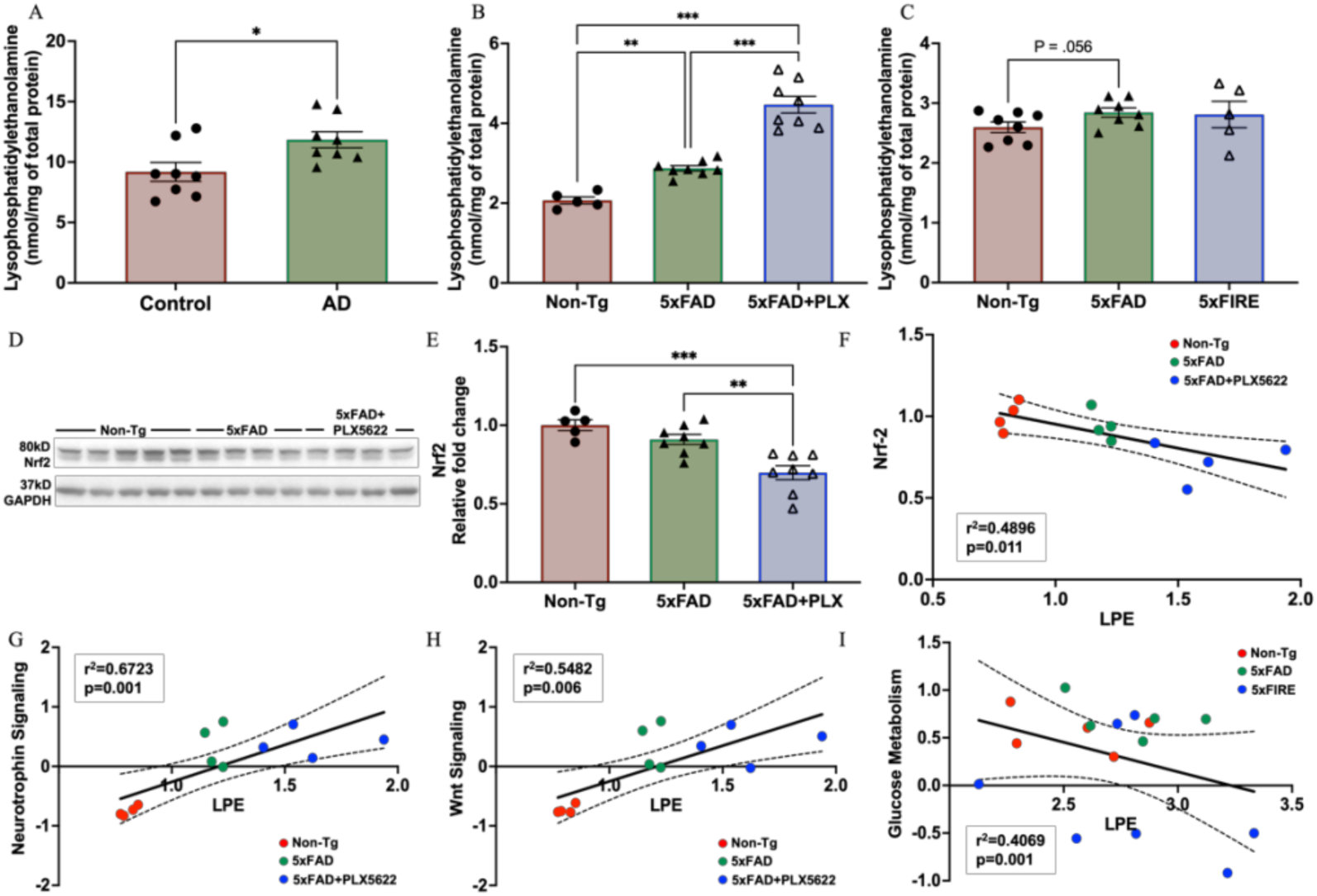
Alzheimer’s disease-associated lysophosphatidylethanolamine accumulation is not restored by microglial elimination. Total LPEs in human AD brains (A) and in pharmacological (B) and genetic (C) microglial-depleted mouse brains. All data presented as mean ± SEM, normalized to WT. Western-blot of Nrf2 (D) and its quantification (E). Two tailed ordinary one-way ANOVA with Turkey correction, *p ≤ 0.05, **p ≤ 0.01, and ***p ≤ 0.001. Correlation analysis among LPE with Nrf2 (F), Neurotrophin signaling score (G), and Wnt signaling score (H) in the pharmacological cohort. (I) Negative correlation between LPE and Glucose metabolism score in the genetic cohort.

Surprisingly, partially eliminating microglia with PLX5622 treatment under amyloidosis exacerbates this accumulation (**Fig. 5B**), particularly non-plasmalogen LPE (**Fig. S4E**), which can be generated following plasmalogen (vinyl-ether bond) cleavage under ROS attack^67,68^. A previous study showed that the 4-HNE-adducted proteins dramatically increased in 5xFAD mice indicating increased oxidative stress and returned to normal levels after blocking the lipid peroxidation^65^. Intriguingly, this trend perfectly aligned with total LPE, but not LPC, levels, suggesting a strong role of oxidative stress in inducing LPE accumulation. Although, in this study, we didn’t find direct evidence of increased oxidative stress contributing to LPE accumulation (**Fig. S5A, S5D, and S5E**), the major antioxidant regulator, Nrf2, was found to decrease after pharmacological microglial elimination (**Fig. 5D-E**). Nrf2 levels also showed a negative correlation with LPE levels (**Fig. 5F**), suggesting there may be a weaker antioxidant response after microglial elimination. Plasmalogen-containing PE/LPE might serve as a potent antioxidant. In contrast to what was observed in the pharmacological cohort, genetic deletion of microglia showed no additional boost of LPE under amyloidosis, except LPE_22:6 (**Fig. S4F**). We attempted to find specific genes or pathways that correlated with LPE levels in both pharmacological and genetic cohorts but could not find any. Despite this, we did find a couple of pathways that correlated strongly with LPE content in the pharmacological cohort only, including neurotrophin and Wnt signaling (**Fig. 5F** and **5G**). On the other hand, LPE levels in the genetic cohort correlated negatively with glucose metabolism (**Fig. 5I**). Although there was no overlap in the negatively correlated genes with LPE observed across both cohorts, the Gene Ontology (GO) terms analysis revealed that such genes were associated with angiogenesis and blood-brain barrier (BBB) regulation within both pharmacological (**Fig. S5F**) and genetic (**Fig. S5G**) cohorts. These findings suggest that increased LPE levels may negatively affect angiogenesis and BBB integrity, which is consistent with a previous study that reported increased angiopathy, BBB dysfunction, and brain hemorrhage using the same genetic microglial depletion mouse model^30^.

### Microglial depletion led to alterations in myelin and energy lipids

Other lipid classes were differentially affected depending on whether microglia were eliminated pharmacologically or genetically. Genetic depletion of microglia under amyloidosis led to a significant reduction of myelin lipids, including cerebroside (**Fig. S6B)** and sulfatide (**Fig. S6D)**, while only mild trends were observed in the pharmacological cohort (**Fig. S6A** and **S6C**). Both cerebroside and sulfatide are prominent structural glycosphingolipids that are found in myelin sheath^69^. Cerebroside, which is crucial for the synthesis and stability of myelin, makes up around 22% of the lipids found in myelin^70^. Sulfatide, the sulfated derivative of cerebroside, has a major role in maintaining the structural integrity of myelin and facilitating connections between axons^71^, and involving gliosis signaling^72^.

Some energy lipid alterations were also observed with microglial depletion. The acyl-carnitine level was found to be significantly increased with pharmacological microglial elimination (**Fig. S6E**), while mitochondria-specific cardiolipin (**Fig. S6G**) and diacylglycerol (**Fig. S6I**) were found to be decreased. On the contrary, the levels of acyl-carnitine (**Fig. S6F**) and diacylglycerol (trending) (**Fig. S6J**) were decreased in the genetic microglial depletion cohort while cardiolipin remained stable (**Fig. S6H**). Diacylglycerol has been shown to regulate mitochondrial function^73^ and ROS signaling^74^ through activating PKC^75^. All these fluctuations can indicate there is an energy metabolism imbalance with microglial depletion. Yet, more detailed characterization is needed to fully understand the distinct metabolism regulation of energy and bioactive lipids with pharmacological and genetic microglial depletion.

## Discussion

Recent studies have intensively investigated the impact of microglia on AD-associated pathologies^26–31^. However, the role of microglia as a potential regulator of brain lipid metabolism remains understudied. Given the number of lipid classes altered at the early stages of AD^9–12^ and the accumulating evidence placing microglia at the forefront of AD pathogenesis, we aimed to assess the effects that microglial depletion has on brain lipid metabolism under AD-like conditions. To this end, we utilized multi-dimensional mass spectrometry-based shotgun lipidomics and revealed specific lipid abnormalities in the AD brain that are either fully driven by microglia or occur independently of these brain-resident macrophages.

BMP is an emerging phospholipid class involved with endo/lysosomal function, which accumulates in a number of lysosomal disorders^76^. A recent study, using a *Grn* knockout mouse to model PGRN loss-of-function mutations that cause frontotemporal dementia, found that PGRN depletion leads to a significant reduction of BMPs^51^. On the other hand, supplementing PGRN rescued BMP losses and prevented the associated lysosomal disorder. In alignment with these findings, our data strongly support a model where increases in BMPs, particularly AA-BMPs, in the context of amyloidosis are associated and driven by increased microglial lysosome (dys)function and increased PGRN levels. Supporting this model, our pharmacological and genetic-based intervention studies demonstrated that partial or complete depletion of microglia is sufficient to prevent amyloidosis-induced AA-BMP accumulation. In addition, we found a positive association between the levels of AA-BMP, the expression of lysosomal genes, protein content of a microglia/macrophage-specific lysosomal marker (CD68), and microglial PGRN RNA and protein levels. Finally, our data also suggest that the loss of microglia-specific PGRN might lead to compensatory mechanisms by other cell types, i.e., neurons based on immunofluorescence results, resulting in non-microglial PGRN upregulation to partially compensate for the loss of microglial lysosomal degradation machinery.

Lysophospholipids have long been considered as markers of neuroinflammation^57–60^. Since microglia are the major drivers of neuroinflammation, we hypothesized microglia would be major generators of lysophospholipids, particularly in a pathological context. Yet, in this study, we surprisingly found that the loss of microglia does not lead to a reduction in either of the two major lysophospholipid classes under either physiological or AD pathological conditions. Upon unbiased correlation analyses, we found that LPC content highly correlates with astrocytic gene expression pathways. These results are still consistent with the established inflammatory role of LPCs, as astrocytes are also known to be major players in neuroinflammatory responses. LPC accumulated under all tested amyloidosis conditions, including in the brains of AD subjects and in the brains of two cohorts of 5xFAD transgenic mice with two different genetic backgrounds. Yet pharmacological and genetic microglial depletion had no impact on LPC levels whatsoever, if anything, LPC levels tended to increase even further. Similarly, microglial elimination also does not ease astrocyte activation, if anything it tends to further exacerbate it based on gene expression and protein levels. These results suggest that LPC accumulation under amyloidosis conditions is driven primarily by astrocyte activation. We could not identify the presumable PLA_2_ responsible for the observed additional increases in LPC content after microglial deletion, as neither cPLA_2_ nor iPLA_2_ levels were increased at the whole cerebrum level. It may be that we could not detect astrocyte-specific increases in either of these PLA_2_s, or that some other PLA_2_ is responsible. In addition, certain lysophospholipid transporters (e.g., Mfsd2a^77^ which transports lysophospholipids into the brain) or lysophospholipid acyltransferases^78^ (converting lysophospholipids back into phospholipids) could also be responsible for these LPC level changes. These hypotheses should be further validated with future single-cell studies or astrocyte-specific manipulation studies.

Microglial depletion, whether induced pharmacologically or genetically, did not reduce the elevated LPE levels in amyloid-bearing brains. On the contrary, PLX5622 further increased LPE content in 5xFAD mice, but did not alter its level in FIRE mice (both in a 5xFAD background). The observed increases in LPE are likely a consequence of either increased oxidative stress or increased PE cleavage by phospholipases. Since increases are most dramatically in PUFA- containing sn-2 acyl LPE, oxidative stress seems to be the most likely explanation^66^. Notably, a recent study demonstrated that PLX5622 has off-target effects on brain vasculature by altering endothelial cholesterol metabolism^80^. As shown in **Fig. S5F**, our pathway analysis of genes negatively correlated with LPE levels identified angiogenesis as the most significant GO term. Interestingly, there appears to be a link between endothelial cholesterol metabolism and angiogenesis, mediated by VEGF^81^. Furthermore, oxidative stress, which inhibits VEGF signaling, is known to regulate angiogenesis^81^. Thus, our study, supported by lipidomics data, suggests that the PLX5622-induced LPE accumulation could be a consequence of its off-target effect on brain vasculature that may induce increases in ROS. On the other hand, upon conducting unbiased correlation analyses between gene expression levels and LPE content, we found a positive correlation with Wnt signaling in the pharmacological cohort, where the strongest effects on LPE levels were observed. Among the genes that included for Wnt signaling provided by NanoString, 16 out of 20 are neuronal-enriched. Wnt signaling is a prominent pathway at the synapse and is required for synaptic plasticity and maintenance in the adult brain, suggesting that LPE increases in the AD brain may also be driven by neurons, potentially by high neuronal oxidative stress. It has been shown that certain levels of oxidative stress can promote canonical Wnt signaling and retinal cell formation and migration^79^. The ROS-associated LPE accumulation and upregulated Wnt signaling observed in this study after microglial elimination might suggest a positive role of PE, particularly plasmalogen PE, in protecting from oxidative stress and maintaining synaptic function. Plasmalogens contain a vinyl-ether bond, which makes them particularly effective as endogenous antioxidants. This correlation agrees with our observation that PLX5622-treated 5xFAD mice had comparable levels of synaptic markers to 5xFAD mice, as previously reported^29^.

The effects of microglia in promoting developmental neurogenesis and myelination by pruning excessive synapse and myelin sheath are well recognized^80,81^. Microglia also have essential roles in clearing myelin debris and maintaining life-long remyelination^52^. However, the impact of microglia in regulating adult myelin homeostasis is underappreciated. As essential components of myelin sheath, changes in myelin lipid levels are likely a sign of impaired myelin integrity. In this study, we found that long-term partial removal of microglia under amyloidosis doesn’t seem to cause myelin impairment since myelin-specific/enriched lipids, i.e., cerebrosides and sulfatides, show no statistical difference after treatment. However, the complete removal of microglia through genetic deletion under amyloidosis conditions led to a significant reduction in myelin lipids. We can exclude the possibility that the loss of myelin lipids is attributed to developmental defects since we did not observe a reduction of myelin lipids at the same age (5 months) in microglia-free (FIRE) non-transgenic mice (data not shown). This suggests that microglia may play an essential role in myelin maintenance/homeostasis, particularly under pathological conditions. In fact, the absence of microglia seems to accelerate the disruption of myelin that would normally occur at later stages in 5xFAD mice. More comprehensive future studies are needed to decipher the relationship between microglia and myelin maintenance.

Lastly, despite a significant amount of overlap and agreement between our pharmacological and genetic mouse cohorts, particularly when comparing lipidomics and RNA profiling data between WT and 5xFAD mouse brains, we did notice that the magnitude of the effects tended to be stronger in the pharmacological cohort. We think this is likely explained by the different genetic backgrounds used. It is well-established that 5xFAD mice develop Aβ pathology faster and more dramatically in the B6SJL background (pharmacological cohort) compared to B6 (genetic cohort) and that females develop a more aggressive pathology than males. In addition, the two cohorts were housed in two different facilities (UTHSCSA and UCI), which means they were exposed to different environments.

## Methods

### Study design and approval

Human Brodmann area 38 samples were provided by the NIH NeuroBioBank (case #481), all 20 samples assessed in this study were obtained from a single site, i.e., the Human Brain and Spinal Fluide Resource Center (Los Angeles, CA). All cases, except for one, were handled by the same neuropathologist (Roscoe Atkinson, MD). Frozen human BA38 samples were kept in a −80°C freezer for storage upon receipt and were lyophilized prior to processing them for lipidomics. Both sexes were included for analysis.

For animal studies, experiments were performed in accordance with the guidelines approved by the Institutional Animal Care and Use Committee (IACUC) of the University of Texas Health San Antonio (UT Health SA) and the University of California, Irvine. No data were excluded, all figures include all samples processed. Animals and other experimental units were assigned randomly to the experimental groups. The sample size for in vivo experiments was based on power calculations and our previous research experience.

### Mice

5xFAD mice (Tg(APPSwFlLon,PSEN1*M146L*L286V)6799Vas/Mmjax), were obtained from the Mutant Mouse Resource and Research Center (MMRRC) at The Jackson Laboratory in two different genetic backgrounds (B6SJL: MMRRC_034840-JAX; and C57BL6: MMRRC_034848-JAX). Mice were housed under 12/12 h light/dark cycles with free access to food and water ad libitum. Only female mice were used for the pharmacological (PLX5622-treated) partial microglial elimination cohort and both male and female mice were used for the genetic (FIRE) full microglial elimination cohort.

For the pharmacological group, 1,200 mg of PLX5622 (free base)/kg was added to OpenStandard Diet (OSD) with 15 kcal% fat (Research diets, INC., New Brunswick, NJ, USA). Control mice were fed with OSD, and experimental mice with OSD+ PLX5622 from 1.5 months of age until 5 months, when tissues were harvested. Csf1r(ΔFIRE/ΔFIRE) mice were generated as previously described^24,30^.

The animals used for this study were all harvested at 5-6 months of age. At the time of collection, animals displayed no apparent health issues. Specifically, they were active, maintained good grooming habits, and had normal weight and activity levels. Prior to sacrifice, the liver, lungs, and heart were inspected, and all appeared healthy.

### Brain preparation

For histological analysis, mice were anesthetized with isoflurane and perfused with PBS 1X for 4 minutes. Left hemibrains were post-fixed in 4% PFA overnight and placed in 10%, 20%, and 30% sucrose solution subsequently, followed by Tissue-Tek® O.C.T. (Sakura Finetek USA, Inc., Torrance, CA, USA) embedding using dry-ice cold isoflurane. Serial 10 µm coronal sections of the brain were collected, the hippocampus was used as a landmark. 2 brain sections from 3 animals per group were quantified (approx. Bregma −2).

Mouse right hemi-cerebra were dissected out, placed in cryotubes, and flash frozen in liquid nitrogen. Frozen brains were lyophilized using benchtop freeze dryer (Labconco, Kansas City, MO, USA) under 0.5 vacuum for 48 hours. Lyophilized brain samples were weighted and transferred to Precellys 0.5 ml tubes pre-filled with small beads. Dried tissues were powdered using Precellys® Evolution Tissue Homogenizer (Bertin technologies, Montigny-le-Bretonneux, France) through one round of manufacture’s “soft program” at 0 °C. Powdered dry cerebrum tissues were weighted and split for lipidomics (∼5 mg), RNA profiling (∼4 mg), and Western blotting (∼10 mg) using disposable anti-static spatulas.

### Lipidomics analysis

Lipidomics analysis was conducted using multi-dimensional mass spectrometry-based shotgun lipidomics (MDMS-SL) following previously described protocols^82^. Briefly, dried and powdered tissues were homogenized in 0.1X PBS. Protein concentrations were measured using bicinchoninic acid (BCA) assay. Lipid extraction from the homogenates, standardized by protein content, employed a modified Bligh and Dyer procedure with added internal standards. Lipid quantification was achieved using triple-quadrupole and orbitrap mass spectrometers (Thermo Fisher Scientific, Waltham, MA, USA), both coupled with a Nanomate device (Advion, Ithaca, NY, USA) and controlled by the Xcalibur system^83^. Data processing, including ion peak selection, baseline correction, data transfer, peak intensity comparison ^13^C deisotoping, and quantitation, was facilitated by a custom-programmed Microsoft Excel macro as previously described^84^, ensuring accurate analysis of lipid molecular species.

### Gene expression analysis

Dried and powdered brain samples underwent RNA extraction using the Maxwell® RSC simplyRNA Tissue Kit. RNA concentration was measured with Qubit RNA BR Assay Kit and RNA Integrity (RIN) was assessed using a TapeStation 4150 and RNA ScreenTape. Subsequently, the multiplex gene expression analysis was conducted using Glial Profiling and Metabolic Pathways Panels and the NanoString nCounter® Technology and nCounter® SPRINT™ Profiler (NanoString Technologies, Seattle, WA, USA). Data processing was performed using the NanoString nSolver 4.0 software, incorporating background reduction based on the average of the Negative Controls, along with standard normalization through Positive Control Normalization and CodeSet Content Normalization techniques. Relative gene expression was calculated after normalization by housekeeping genes.

### Western blotting

Dried and powdered brain samples were homogenized in 1X NP40 lysis buffer with Halt Protease and Phosphatase Inhibitor Cocktails (Thermo Fisher Scientific, Waltham, MA, USA) using Precellys® Evolution Tissue Homogenizer (Bertin technologies, Montigny-le-Bretonneux, France). Homogenates were centrifuged at 12,000 g for 30 min at 4 °C; protein concentration of supernatants was determined using the Bio-Rad protein assay (Bio-Rad, Hercules, CA, USA). Supernatants were run with NuPage 4–12% Bis-Tris gels (Life Technologies, Grand Island, NY, USA) under reducing conditions. PageRuler Plus Prestained Protein Ladder (Thermo Fisher Scientific, Waltham, MA, USA) was used as indicator. PVDF membranes (CliniSciences, Nanterre, France) with the transferred protein were incubated with primary antibodies (1:1000-2000 dilutions) of anti-6E10 (mouse, BioLegend, Inc., San Diego, California, USA), anti-Homer1 (rabbit, Cell Signaling Technology, Boston, MA, USA), anti-CD68 (E307V) (rabbit, Cell Signaling Technology, Boston, MA, USA), anti-LAMP-1 (1D4B) (rat, Thermo Fisher Scientific, Waltham, MA, USA); anti-4HNE (12F7) (mouse, Thermo Fisher Scientific, Waltham, MA, USA), anti-PGRN (sheep, R&D Systems, Minneapolis, MN, USA), anti-Phospho-PLA2G4A (Ser505) (Rabbit, Proteintech Group, Inc, Rosemont, IL, USA), anti-iPLA2 (D-4) (mouse, Santa Cruz Biotechnology, Dallas, TX, USA), anti-HO-1 (rabbit, Proteintech Group, Inc, Rosemont, IL, USA), anti-Nrf2 (rabbit, Abcam, Waltham, MA, USA), anti-GFAP (rabbit, Thermo Fisher Scientific, Waltham, MA, USA), anti-GAPDH (rabbit, Cell Signaling Technology, USA) overnight at 4 °C, followed by horseradish peroxidase (HRP)-linked secondary antibodies (Cell Signaling Technology, Boston, MA, USA) for 1 h at room temperature.

Pierce™ ECL Western Blotting Substrate was used for protein detection followed by autoradiography film exposure (HyBlot CL). A subset of these PVDF membranes underwent a reblotting process with stripping buffer (Thermo Fisher Scientific, Waltham, MA, USA). Protein expression levels were quantified utilizing ImageJ software normalized relative to GAPDH expression.

### Immunofluorescence staining

Frozen brain slices were blocked using 10% normal goat serum (Sigma, USA) for 1 h at room temperature, then incubated with the following primary antibodies: anti-Iba1 (rabbit, FUJIFILM Cellular Dynamics, Inc., Madison, WI, USA), anti-PGRN (sheep, R&D Systems, Inc., Minneapolis, MN, USA), anti-MOAB-2 (mouse, MilliporeSigma, Burlington, MA, USA) at 4 °C overnight, washed three times, then incubated with fluorescence-labeled Alexa Fluor secondary antibodies (Invitrogen, USA) 1 h at room temperature; washed three times, and mounted with DAPI. Images were captured with a confocal laser-scanning microscope (Zeiss LSM710, USA) or Fluorescence Microscope BZ-X800 (KEYENCE, Japan).

### Statistics

Data in the figures are presented as mean ± SEM. All the statistical analyses for gene transcript counts from Nanostring were performed using the NanoString nSolver recommended test (heteroscedastic Welch’s t-Test). All other statistical analyses were performed using Prism (GraphPad) and MetaboAnalyst (https://www.metaboanalyst.ca/). Specific tests and multiple comparison corrections performed are specified in the figure legends.

## Acknowledgements

Human brain tissue specimens were obtained from the Human Brain and Spinal Fluid Resource Center located at the VA Greater Los Angeles Healthcare System, West Los Angeles Healthcare Center, Los Angeles, CA 90073. This center is sponsored of Mental Health (NIMH), the National multiple Sclerosis Society, and the Department of Veterans Affairs.

## Data availability

The authors declare that the data supporting the findings of this study are available within the paper and its Supplementary Information files. Should any raw data files be needed in another format they are available from the corresponding author upon reasonable request.

---

**Supplementary Figure 1.**
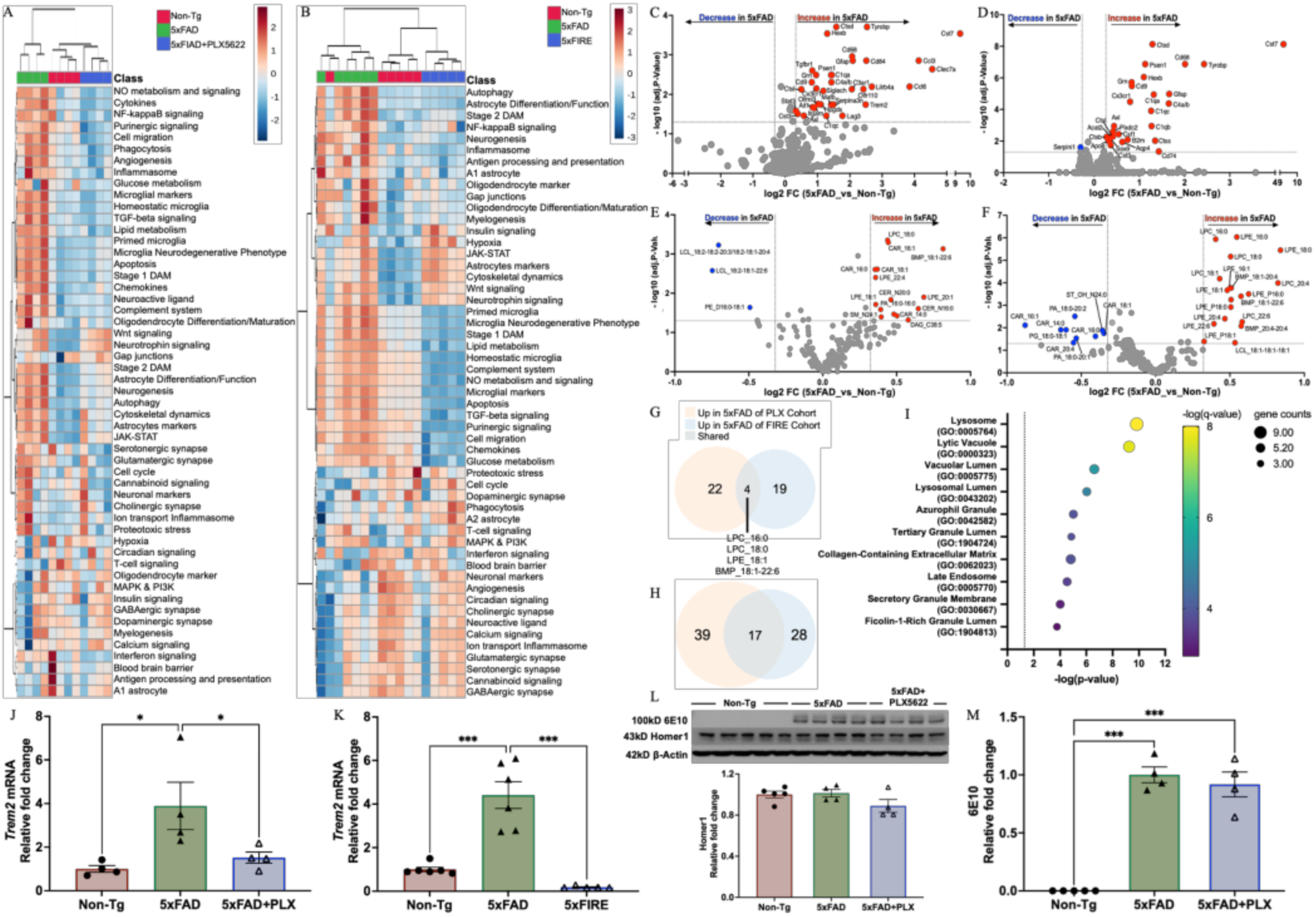
Overall brain lipidome and transcriptome patterns under amyloidosis. Heatmaps of transcriptomic pathways in pharmacological (A) and genetic (B) microglia deficient cohorts. Volcano plots showing the significantly changed genes under amyloidosis in pharmacological (C) and genetic (D) cohorts. Volcano plots showing the significantly changed lipid species under amyloidosis in pharmacological (E) and genetic (F) cohorts. Venn diagrams show increased lipid species (G) and genes (H) shared in 5xFAD mice of both pharmacological and genetic cohorts. Data transformation: log10; data scaling: mean. Adjusted p≤ 0.05, fold-change ≥ 0.25. Bubble plot showing top 10 Gene Ontology (GO) terms analysis (I) of shared genes. The q-value is an adjusted p-value calculated using the Benjamini-Hochberg method for correction. The Trem2 mRNA levels in both pharmacological (J) and genetic (K) cohorts. (L) APP level measured by 6E10 and Homer1 westerns (up) and Homer1 quantification (down). (M) The quantification of 100kD 6E10. Ordinary one-way ANOVA with Turkey correction, *p ≤ 0.05, **p ≤ 0.01, ***p ≤ 0.001.

**Supplementary Figure 2.**
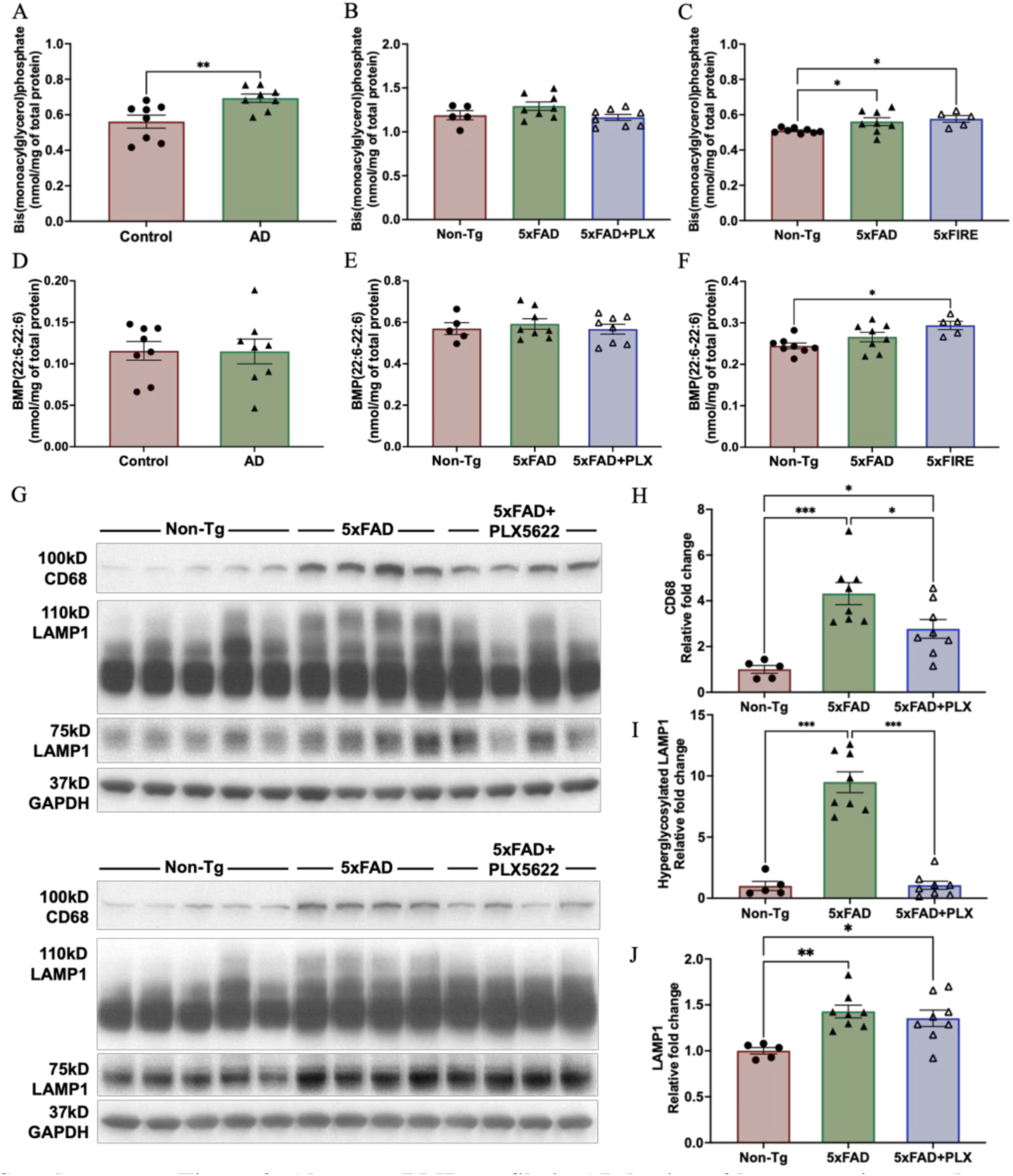
Aberrant BMP profile in AD brains of human patients and mouse models. Total BMP changes in human AD brains (A), pharmacological (B) and genetic (C) microglia-deficiency conditions. Docosahexaenoic acid (DHA)-specific BMP (DHA-BMP) changes in human AD brains (D), pharmacological (E) and genetic (F) microglia-deficiency conditions. Western-blot showing CD68 and LAMP1 (G) in pharmacological microglia-deficiency cohort. Quantification of CD68 (H), normal LAMP1 (75kD, in I) and hyperglycosylated LAMP1 (110kD, in J) in pharmacological microglia-deficiency cohort. All data presented as mean ± SEM, normalized to WT. Ordinary one-way ANOVA with Turkey correction, *p ≤ 0.05, **p ≤ 0.01, ***p ≤ 0.001.

**Supplementary Figure 3.**
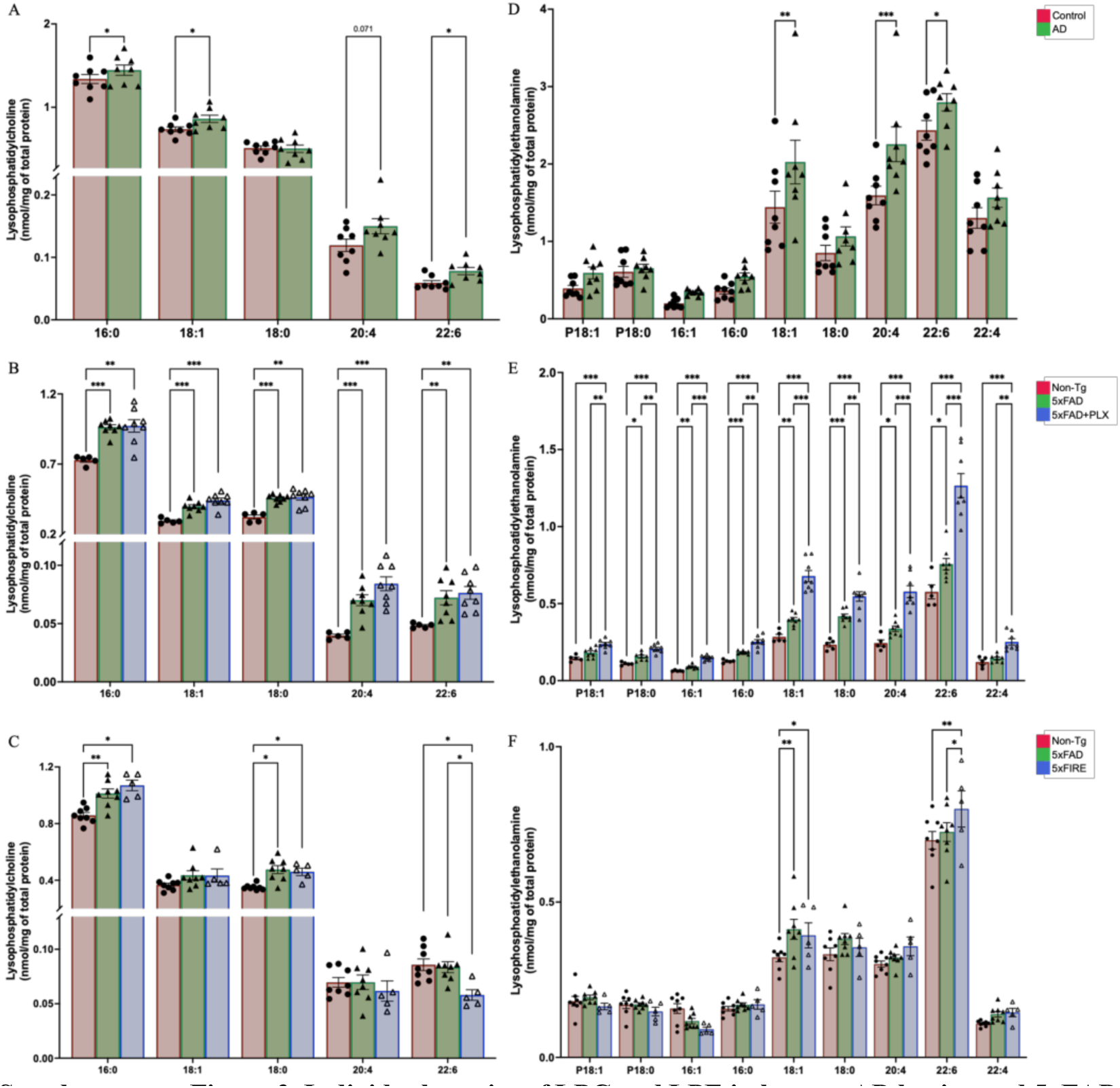
Individual species of LPC and LPE in human AD brains and 5xFAD mice with pharmacological and genetic microglial eliminations. LPC species in human AD brains (A), pharmacological (B) and genetic (C) cohorts. LPE species in human AD brains (D), pharmacological (E) and genetic (F) cohorts. Only showing the species that shared with human. All data presented as mean ± SEM. Multiple unpaired t-test with Holm-Šídák correction, *p ≤ 0.05, **p ≤ 0.01, ***p ≤ 0.001.

**Supplementary Figure 4.**
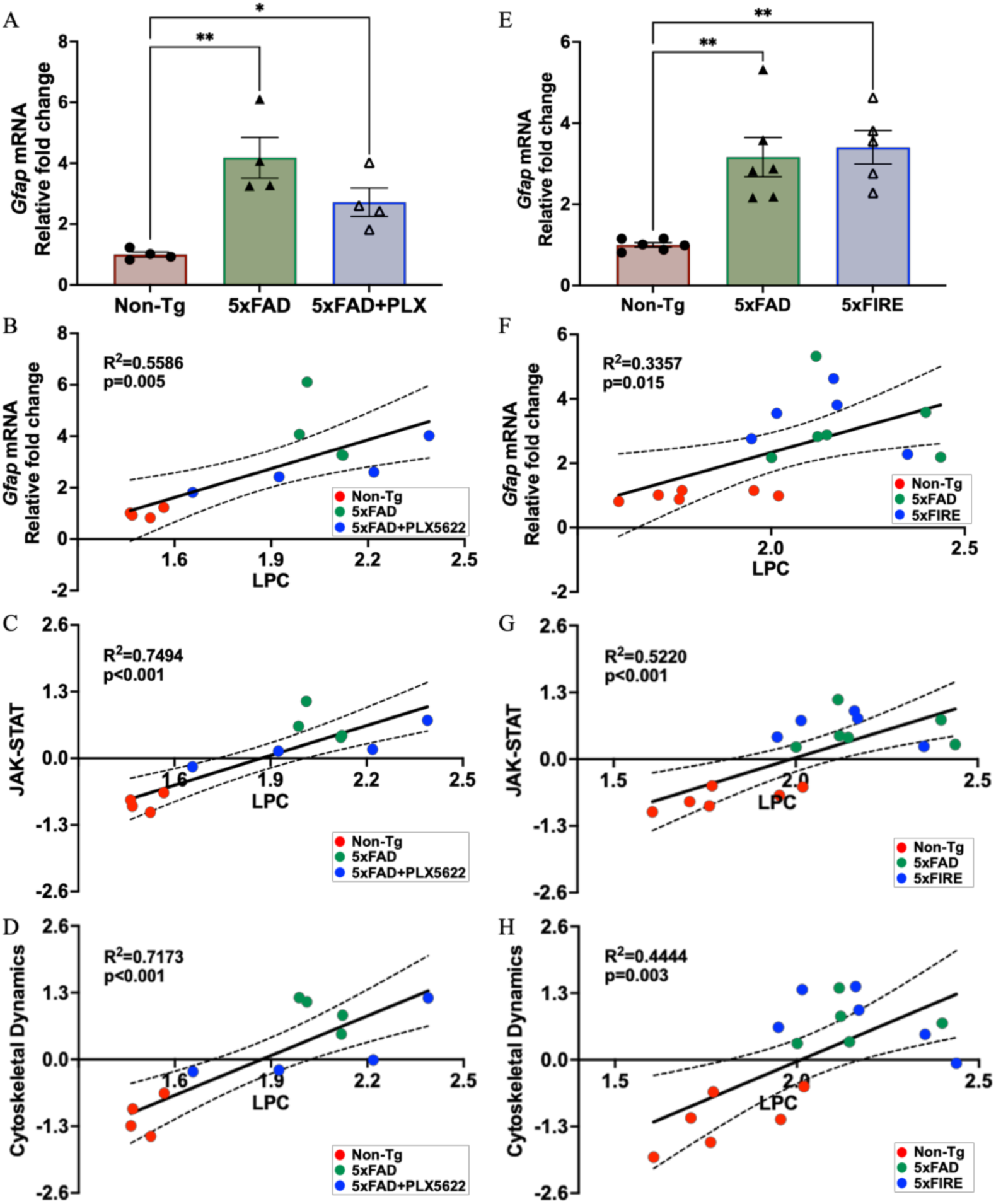
Correlations between LPC and transcriptomic pathways in pharmacological microglial elimination cohort. *Gfap* mRNA levels in the pharmacological (A) and genetic (E) cohorts. Correlations between LPC and *Gfap* mRNA levels in the pharmacological (B) and genetic (F) cohorts. Correlations between LPC and cytoskeletal dynamics in the pharmacological (C) and genetic (G) cohorts. (D) Correlations between LPC and astrocyte markers in the pharmacological (D) and genetic (H) cohorts.

**Supplementary Figure 5.**
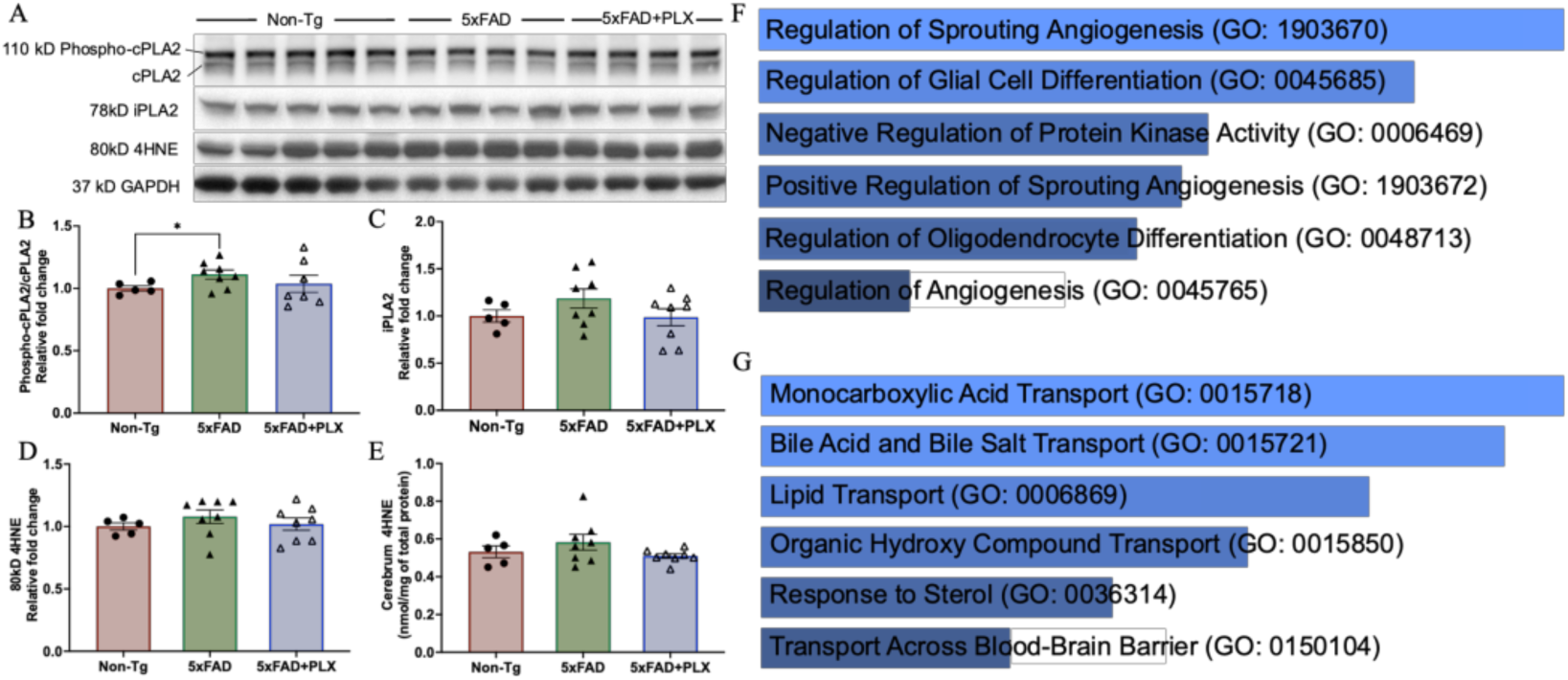
Metabolism signaling underlying lysospholipids alteration in amyloidosis and pharmacological microglial elimination cohorts. Western-blot analysis (A) of calcium-dependent and independent phospholipase A2 (phosphor-cPLA2 and iPLA2, respectively), 4HNE levels and their quantification results (B, C, and D). (E) 4-HNE level measure by MDMS-shotgun lipidomics. Gene Ontology (GO) terms analysis of genes that negatively correlated with LPE in pharmacological (F) and genetic (G) microglial elimination cohort. Data transformation: square root for the pharmacological cohort, cube root for the genetic cohort; data scaling: pareto for the pharmacological cohort, mean for the genetic cohort. All data presented as mean ± SEM, normalized to WT. Two tailed two-way ANOVA with Turkey correction, *p ≤ 0.05, **p ≤ 0.01, ***p ≤ 0.001.

**Supplementary Figure 6.**
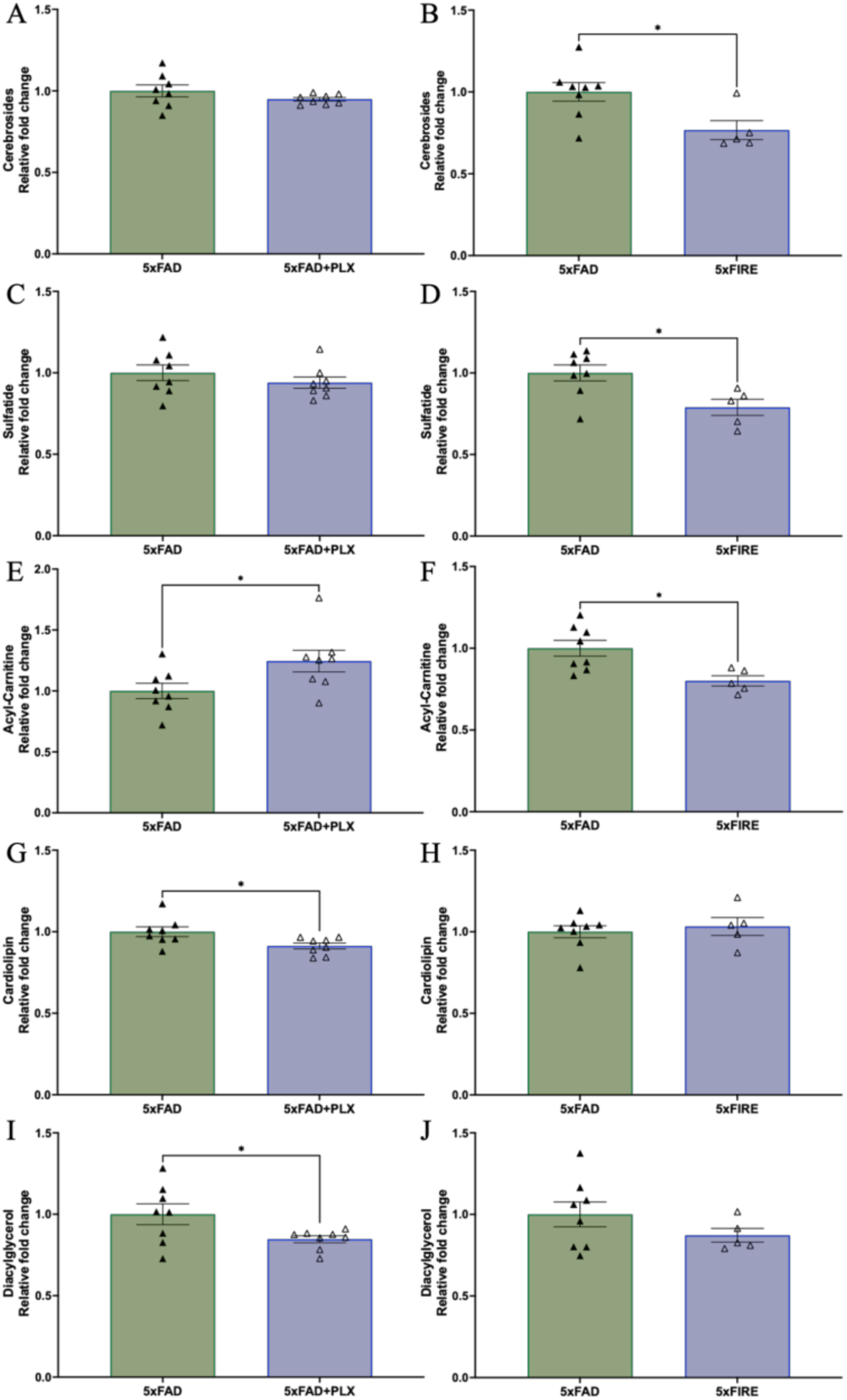
Additional lipid classes change with pharmacological and genetic microglial eliminations under amyloidosis and WT backgrounds. Relative changes of cerebrosides (A), sulfatide (C), carnitine (E), cardiolipin (G), and diacylglycerol (I) in pharmacological microglial elimination under amyloidosis. Relative changes of cerebrosides (B), sulfatide (D), carnitine (F), cardiolipin (H), and diacylglycerol (J) in genetic microglial elimination under amyloidosis. All data presented as mean ± SEM, normalized to WT. Ordinary one-way ANOVA with Turkey correction, *p ≤ 0.05, **p ≤ 0.01, ***p ≤ 0.001.

**Supplementary Table 1.**
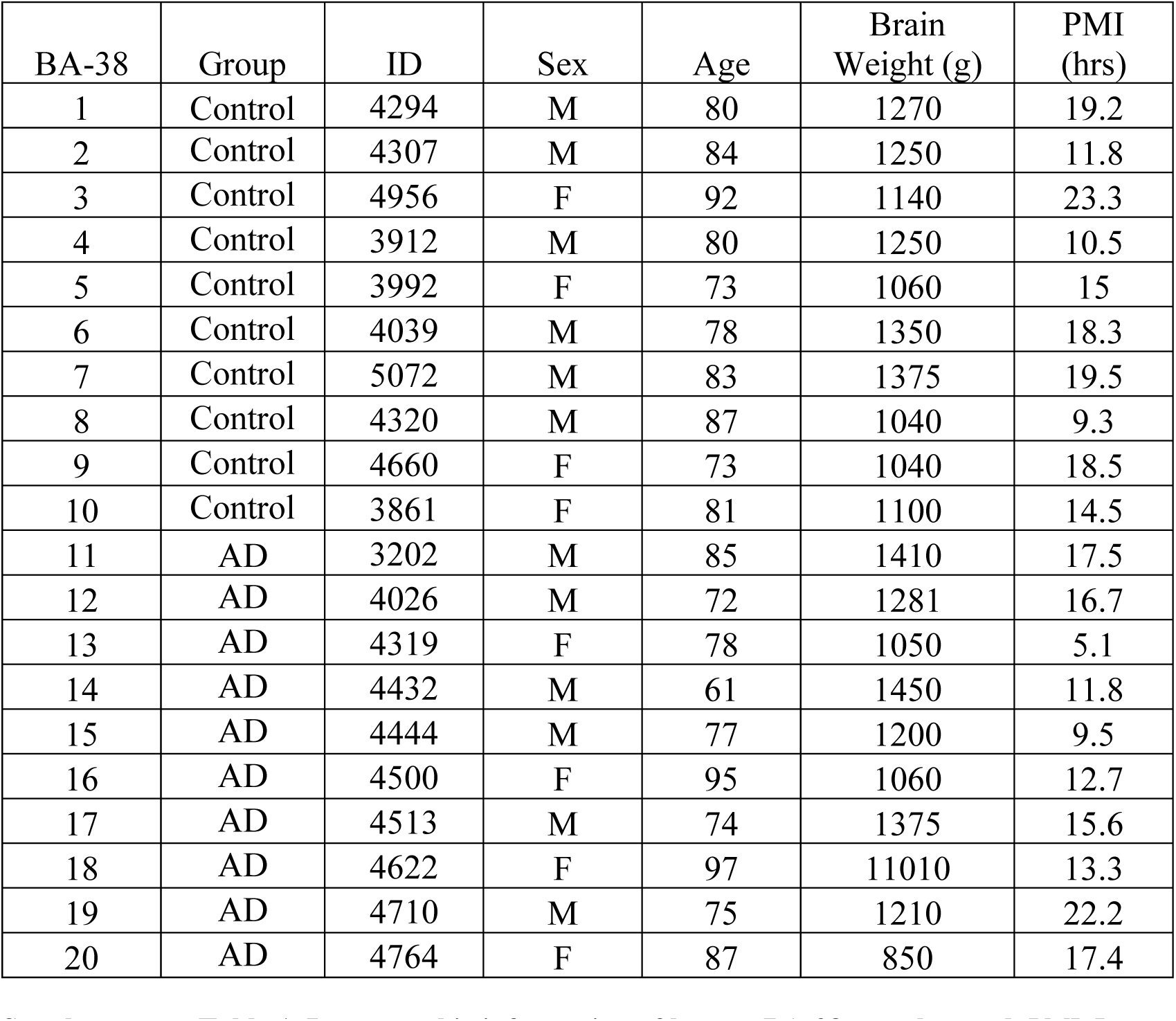
Demographic information of human BA-38 samples used. PMI, Post mortem interval.

## Notes

### Competing Interest Statement

The authors have declared no competing interest.

