## supplementary figures for "Microglia-Dependent and Independent Modulation of Brain Lipid Metabolism in Alzheimer’s Disease Revealed by Pharmacological and Genetic Microglial Depletion"

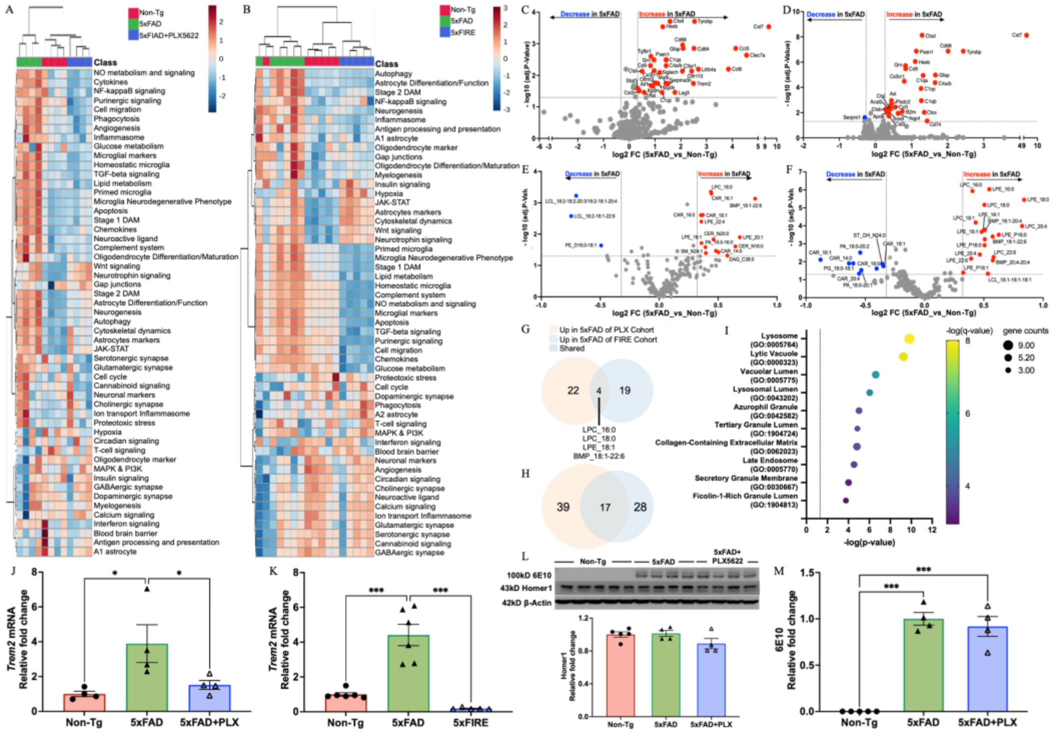

**Supplementary Figure 1. Overall brain lipidome and transcriptome patterns under amyloidosis.** Heatmaps of transcriptomic pathways in pharmacological (A) and genetic (B) microglia deficient cohorts. Volcano plots showing the significantly changed genes under amyloidosis in pharmacological (C) and genetic (D) cohorts. Volcano plots showing the significantly changed lipid species under amyloidosis in pharmacological (E) and genetic (F) cohorts. Venn diagrams show increased lipid species (G) and genes (H) shared in 5xFAD mice of both pharmacological and genetic cohorts. Data transformation: log10; data scaling: mean. Adjusted p≤ 0.05, fold-change ≥ 0.25. Bubble plot showing top 10 Gene Ontology (GO) terms analysis (I) of shared genes. The q-value is an adjusted p-value calculated using the Benjamini-Hochberg method for correction. The Trem2 mRNA levels in both pharmacological (J) and genetic (K) cohorts. (L) APP level measured by 6E10 and Homer1 westerns (up) and Homer1 quantification (down). (M) The quantification of 100kD 6E10. Ordinary one-way ANOVA with Turkey correction, $*p \leq 0.05$, $**p \leq 0.01$, $***p \leq 0.001$.

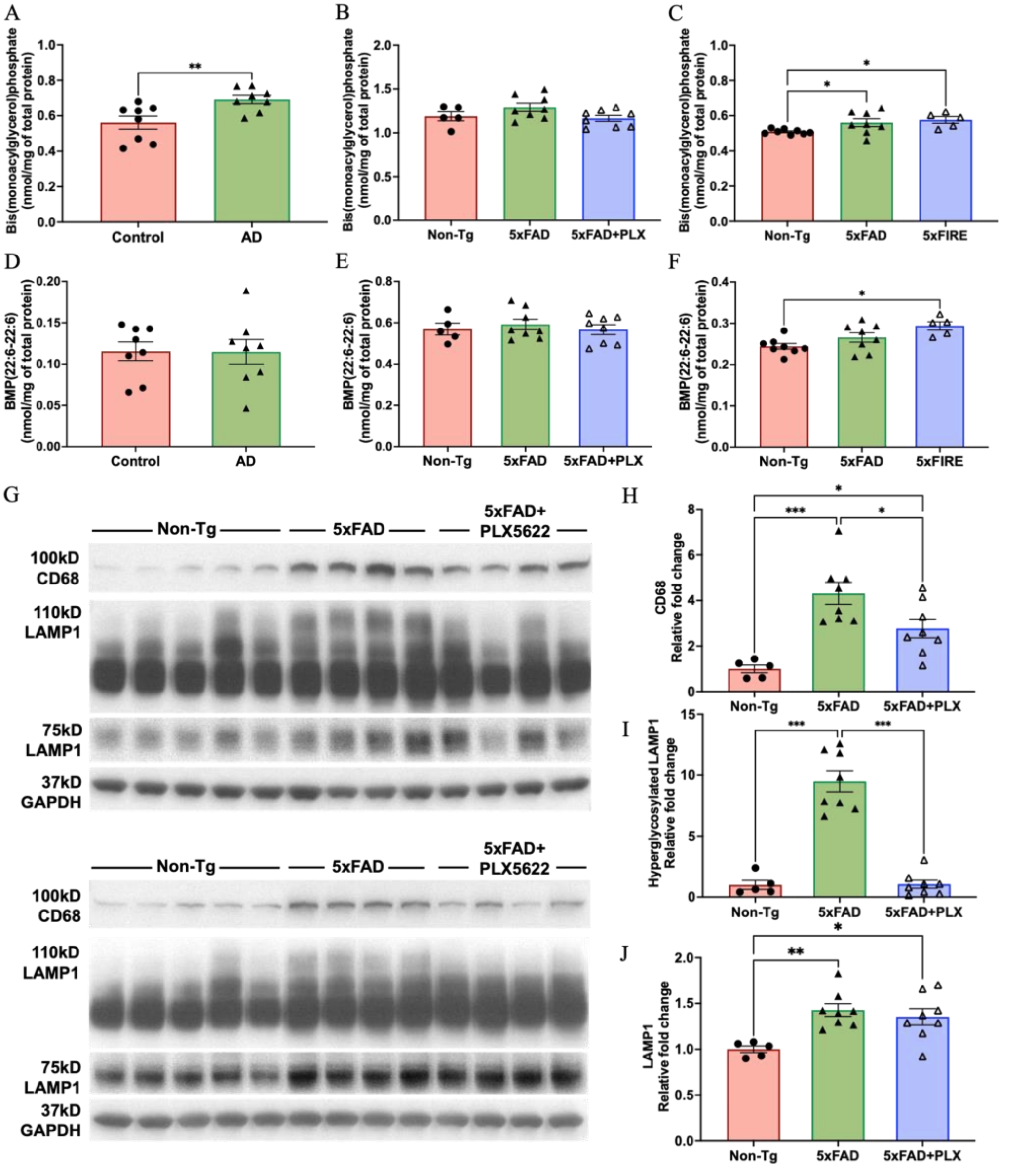


**Supplementary Figure 2. Aberrant BMP profile in AD brains of human patients and mouse models.** Total BMP changes in human AD brains (A), pharmacological (B) and genetic (C) microglia-deficiency conditions. Docosahexaenoic acid (DHA)-specific BMP (DHA-BMP) changes in human AD brains (D), pharmacological (E) and genetic (F) microglia-deficiency conditions. Western-blot showing CD68 and LAMP1 (G) in pharmacological microglia-deficiency cohort. Quantification of CD68 (H), normal LAMP1 (75kD, in I) and hyperglycosylated LAMP1 (110kD, in J) in pharmacological microglia-deficiency cohort. All data presented as mean ± SEM, normalized to WT. Ordinary one-way ANOVA with Turkey correction, $*p \le 0.05$, $**p \le 0.01$, $***p \le 0.001$.

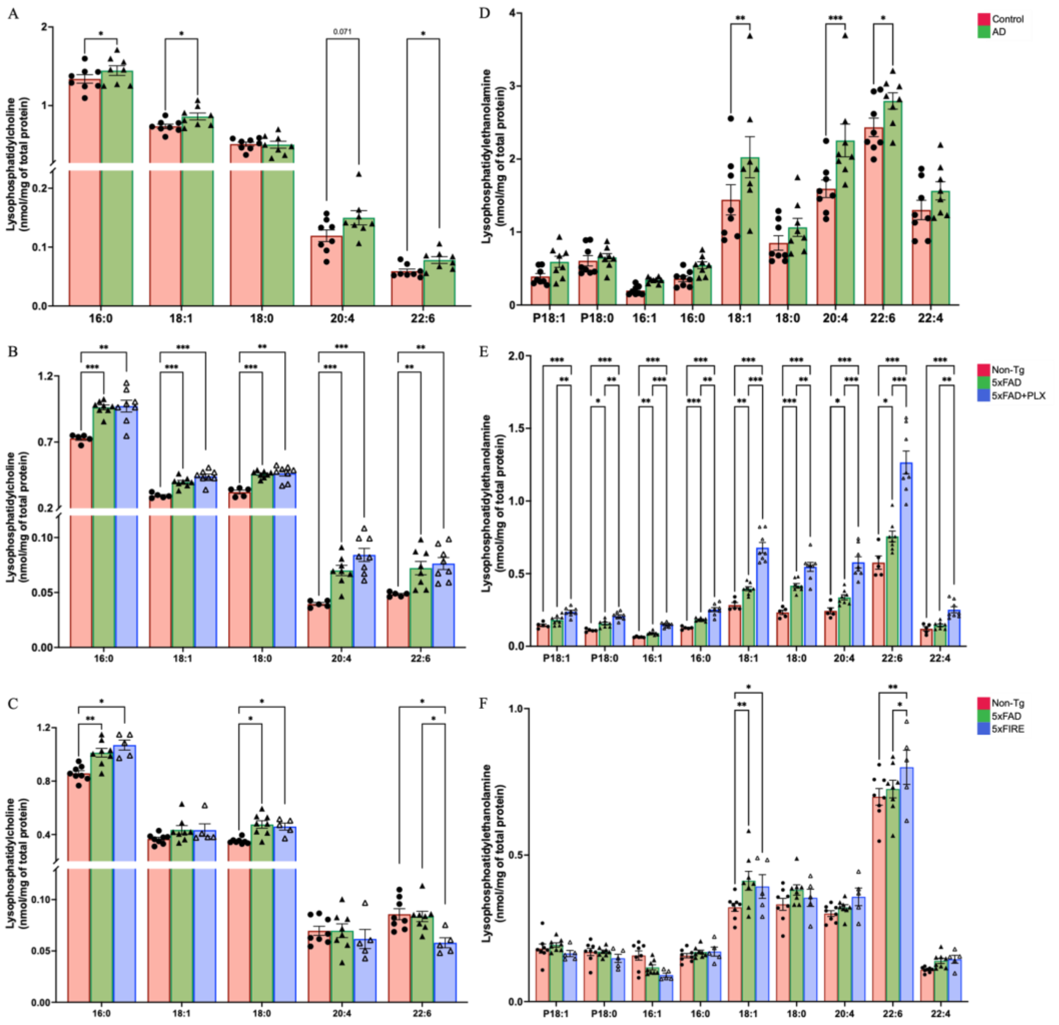


**Supplementary Figure 3. Individual species of LPC and LPE in human AD brains and 5xFAD mice with pharmacological and genetic microglial eliminations.** LPC species in human AD brains (A), pharmacological (B) and genetic (C) cohorts. LPE species in human AD brains (D), pharmacological (E) and genetic (F) cohorts. Only showing the species that shared with human. All data presented as mean ± SEM. Multiple unpaired t-test with Holm-Šídák correction, $*p \leq 0.05$, $**p \leq 0.01$, $***p \leq 0.001$.

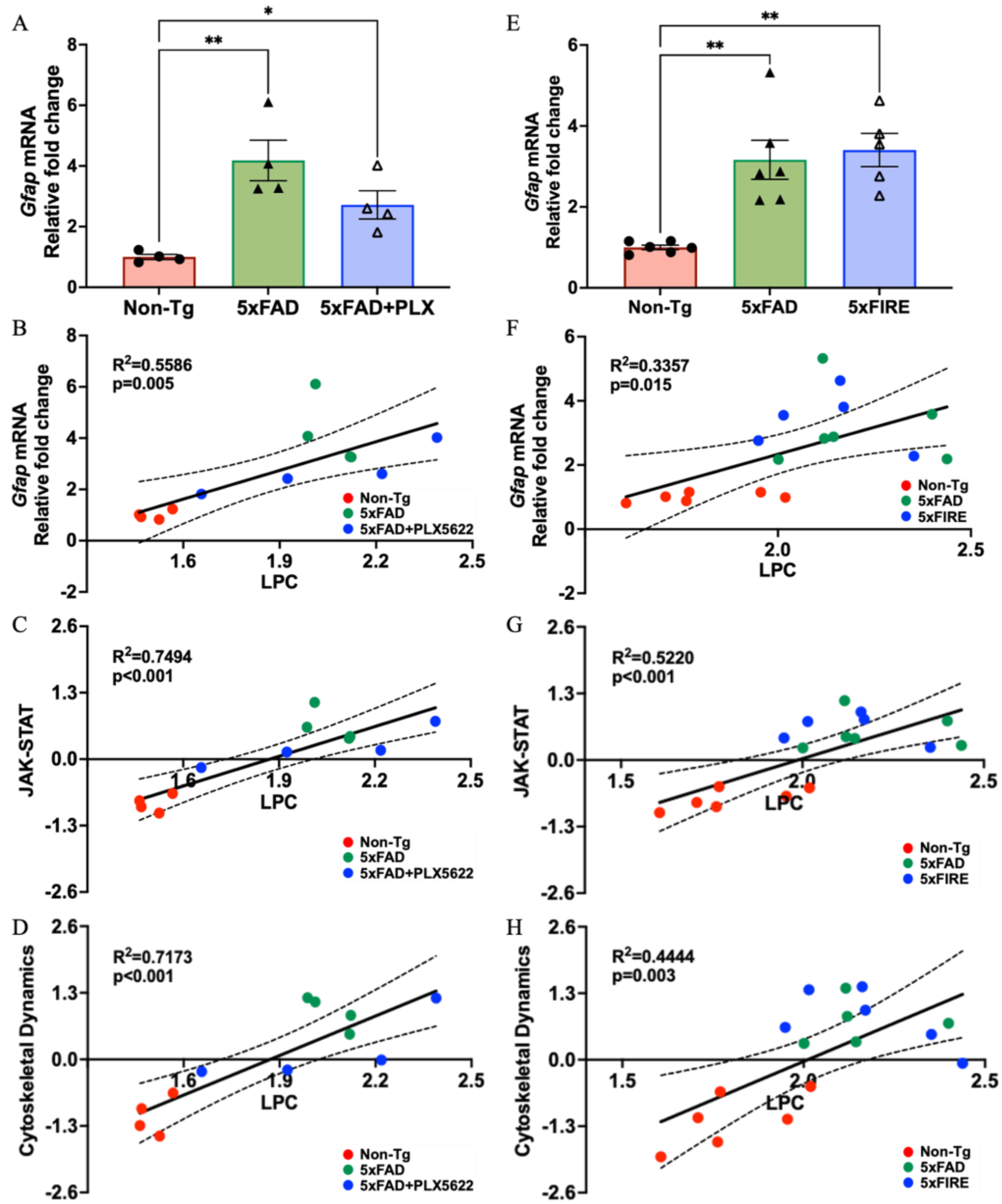


**Supplementary Figure 4. Correlations between LPC and transcriptomic pathways in pharmacological microglial elimination cohort.** *Gfap* mRNA levels in the pharmacological (A) and genetic (E) cohorts. Correlations between LPC and *Gfap* mRNA levels in the pharmacological (B) and genetic (F) cohorts. Correlations between LPC and cytoskeletal dynamics in the pharmacological (C) and genetic (G) cohorts. (D) Correlations between LPC and astrocyte markers in the pharmacological (D) and genetic (H) cohorts.

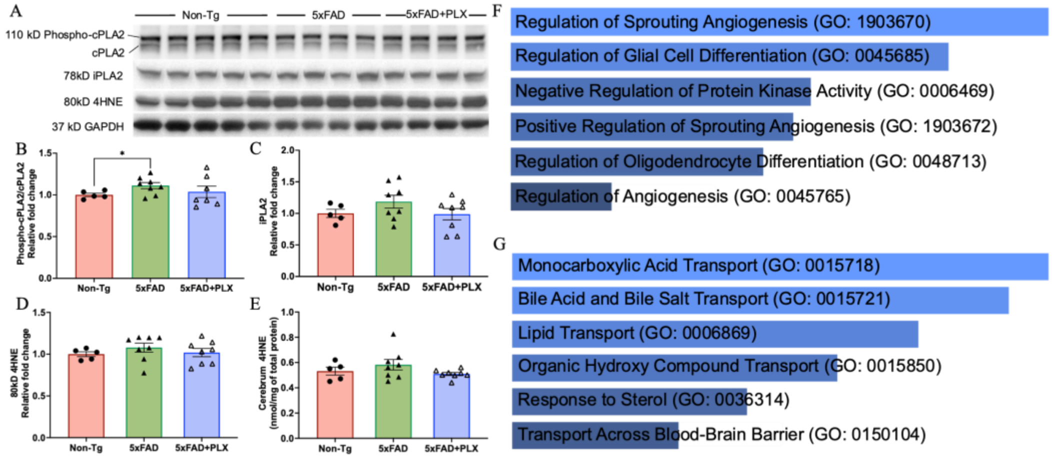


**Supplementary Figure 5. Metabolism signaling underlying lysospholipids alteration in amyloidosis and pharmacological microglial elimination cohorts.** Western-blot analysis (A) of calcium-dependent and independent phospholipase A2 (phosphor-cPLA2 and iPLA2, respectively), 4HNE levels and their quantification results (B, C, and D). (E) 4-HNE level measure by MDMS-shotgun lipidomics. Gene Ontology (GO) terms analysis of genes that negatively correlated with LPE in pharmacological (F) and genetic (G) microglial elimination cohort. Data transformation: square root for the pharmacological cohort, cube root for the genetic cohort; data scaling: pareto for the pharmacological cohort, mean for the genetic cohort. All data presented as mean ± SEM, normalized to WT. Two tailed two-way ANOVA with Turkey correction, $*p \leq 0.05$, $**p \leq 0.01$, $***p \leq 0.001$.

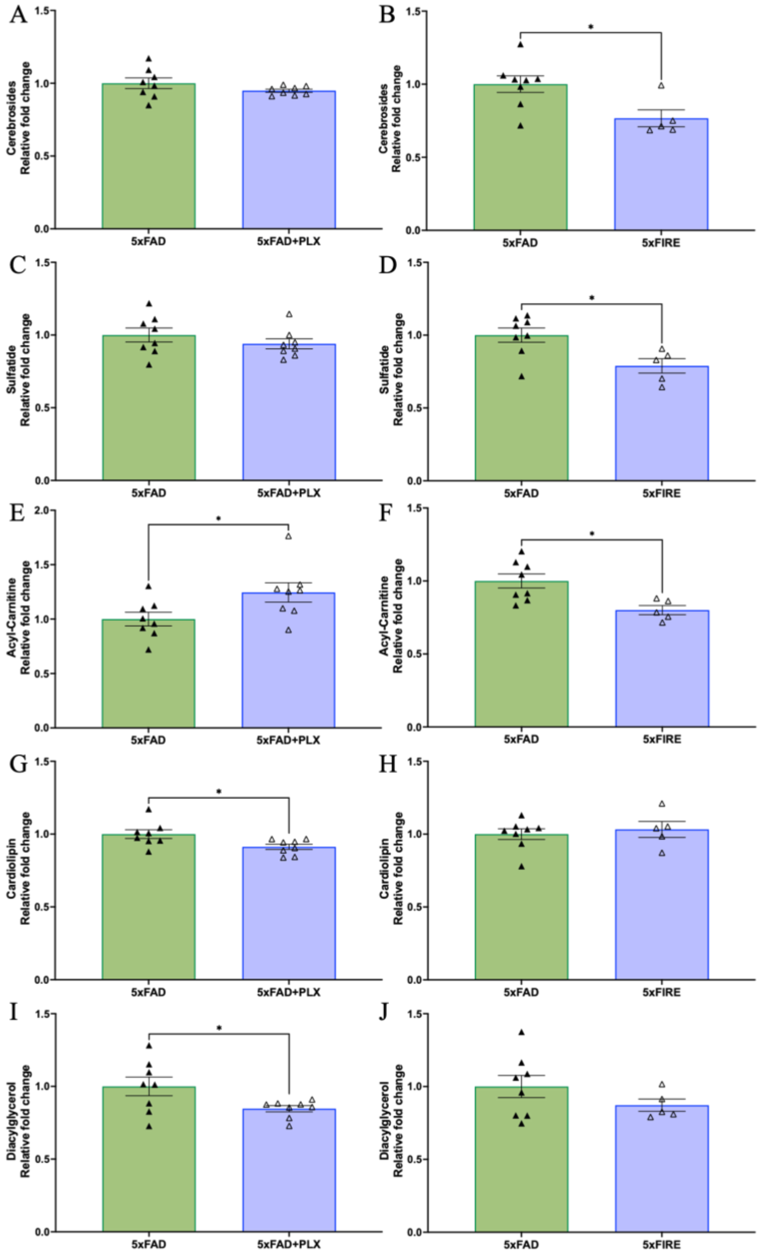


**Supplementary Figure 6. Additional lipid classes change with pharmacological and genetic microglial eliminations under amyloidosis and WT backgrounds.** Relative changes of cerebrosides (A), sulfatide (C), carnitine (E), cardiolipin (G), and diacylglycerol (I) in pharmacological microglial elimination under amyloidosis. Relative changes of cerebrosides (B), sulfatide (D), carnitine (F), cardiolipin (H), and diacylglycerol (J) in genetic microglial elimination under amyloidosis. All data presented as mean ± SEM, normalized to WT. Ordinary one-way ANOVA with Turkey correction, $*p \leq 0.05$, $**p \leq 0.01$, $***p \leq 0.001$.

| BA-38 | Group | ID | Sex | Age | Brain Weight (g) | PMI (hrs) |
|---|---|---|---|---|---|---|
| 1 | Control | 4294 | M | 80 | 1270 | 19.2 |
| 2 | Control | 4307 | M | 84 | 1250 | 11.8 |
| 3 | Control | 4956 | F | 92 | 1140 | 23.3 |
| 4 | Control | 3912 | M | 80 | 1250 | 10.5 |
| 5 | Control | 3992 | F | 73 | 1060 | 15 |
| 6 | Control | 4039 | M | 78 | 1350 | 18.3 |
| 7 | Control | 5072 | M | 83 | 1375 | 19.5 |
| 8 | Control | 4320 | M | 87 | 1040 | 9.3 |
| 9 | Control | 4660 | F | 73 | 1040 | 18.5 |
| 10 | Control | 3861 | F | 81 | 1100 | 14.5 |
| 11 | AD | 3202 | M | 85 | 1410 | 17.5 |
| 12 | AD | 4026 | M | 72 | 1281 | 16.7 |
| 13 | AD | 4319 | F | 78 | 1050 | 5.1 |
| 14 | AD | 4432 | M | 61 | 1450 | 11.8 |
| 15 | AD | 4444 | M | 77 | 1200 | 9.5 |
| 16 | AD | 4500 | F | 95 | 1060 | 12.7 |
| 17 | AD | 4513 | M | 74 | 1375 | 15.6 |
| 18 | AD | 4622 | F | 97 | 11010 | 13.3 |
| 19 | AD | 4710 | M | 75 | 1210 | 22.2 |
| 20 | AD | 4764 | F | 87 | 850 | 17.4 |

**Supplementary Table 1. Demographic information of human BA-38 samples used.** PMI, Post mortem interval.
